# Synthesis and Preclinical Development of a Novel ^68^Ga/^89^Zr-Labelled ανβ6-Integrin Targeting Trimer

**DOI:** 10.1101/2025.11.05.686720

**Authors:** Giacomo Gariglio, Fernando A. Patiño Álvarez, Maximilian A. Zierke, Stefan Stangl, Tim Rheinfrank, Nadine Holzleitner, Susanne Kossatz, Clemens Decristoforo

## Abstract

The αvβ6 integrin has emerged as a valuable target for theranostic applications in nuclear medicine with high applicability across a variety of cancers, including head-and-neck, lung, breast, and pancreatic carcinomas. [⁶⁸Ga]Ga-Trivehexin is a prominent example of a diagnostic tracer targeting this integrin. In this work, we aimed to expand on this concept by developing FSC(PEG4-αvβ6)₃, a novel tracer that retains the Trivehexin design, but features PEGylated spacers and replaces the TRAP chelator with Fusarinine C (FSC), enabling labelling with Zirconium-89 in addition to Gallium-68. Preclinical characterization of [⁶⁸Ga]Ga/[⁸⁹Zr]Zr-FSC(PEG4-αvβ6)₃ included affinity determination towards the αvβ6 integrin and cellular uptake studies in αvβ6-positive H2009 cells. A subcutaneously xenografted H2009 tumor model was used to assess the PET imaging potential and biodistribution at early time points with the Gallium-68-labelled compound, and at later time points (up to 6 days post-injection) with the Zirconium-89-labelled version. While [⁶⁸Ga]Ga-FSC(PEG4-αvβ6)₃ exhibited moderate binding to αvβ6, its affinity, cellular internalization, and tumor uptake *in vivo* were lower compared to [⁶⁸Ga]Ga-Trivehexin. Notably, this decreased target engagement was associated with reduced nonspecific binding, which we primarily attributed to the incorporation of PEGylated linkers. Despite indication of *in vivo* degradation of [⁸⁹Zr]Zr-FSC(PEG4-αvβ6)₃, still a meaningful evaluation of pharmacokinetics and biodistribution at extended time points was feasible, indicating its suitability for prolonged imaging studies.

## INTRODUCTION

Integrins are a large family of transmembrane cell-surface glycoproteins that mediate key biological processes, most notably bidirectional signal transduction across the cell membrane and cellular adhesion through linkage of the cytoskeleton to the extracellular matrix^1, 2^. The integrin repertoire consists of 18 α and 8 β subunits, which non-covalently associate in various combinations to form 24 distinct heterodimeric receptors, each characterized by specific ligand-binding properties. RGD-binding integrins are a subset of 8 integrin receptors that specifically recognize the tripeptide sequence Arg-Gly-Asp (RGD), a common motif present in several extracellular matrix (ECM) proteins, including fibronectin and vitronectin. Upon ligand binding, these integrins undergo clustering in focal adhesion complexes, thereby initiating signal transduction pathways critical for various cellular processes as changes in shape, migration, proliferation and survival ^2^.

The integrin dimer αvβ6 primarily activates TGF-β, which normally inhibits epithelial cell proliferation to maintain tissue homeostasis, but in cancer cells this growth-suppressive effect is lost, instead it is restricted to the tumor microenvironment. There, the excessive activated TGF-β promotes immunosuppression, angiogenesis, and tumor invasiveness ^3–6^. In this context, αvβ6 integrin promotes the infiltrative growth of many kinds of malignant epithelial tumors. Upregulation of αvβ6 has been reported in oral squamous cell carcinoma (OSCC)^7–9^, breast carcinoma^10, 11^, gastric carcinoma^12, 13^, pancreatic ductal adenocarcinoma (PDAC)^14, 15^, colorectal carcinoma (CRC)^16–18^, cholangiocarcinoma^19, 20^, non-small cell lung cancer (NSCLC)^21^ and ovarian cancer^22, 23^. Importantly, elevated expression levels have been associated with increased tumor invasiveness, which clinically correlates with metastatic progression and poorer patient survival^18, 24–25^. Beyond oncology, its clinical scope extends also to fibrotic diseases such as idiopathic pulmonary fibrosis (IPF) and potentially even COVID-19–related syndromes^26^. Owing to its absence in most healthy adult tissues, its widespread overexpression on the cell surface across these tumor types, along with the rapid internalization of the ligand-receptor complex within 30–60 minutes^27^, αvβ6 integrin has emerged as a highly promising target for innovative diagnostic and therapeutic strategies^28^.

Since the early 2000s, several αvβ6-targeted probes for nuclear medicine applications have been developed^29–37^, with some already evaluated in clinical trials^38–41^. This work has predominantly focused on peptide-based agents, given their synthetic accessibility, tunable structure, low molecular weight, minimal immunogenicity, and rapid clearance from normal tissues^42,43^.

To optimize the target binding, a library of ligand candidates has been generated utilizing strategies such as 1-bead-1-compound combinatorial libraries,as well as phage- and yeast-display, with additional adaptation of naturally occurring protein fragments. Subsequent screening identified several candidates of 7–20 residues, revealing that the minimal RG/TDLXXL motif (X = any α-amino acid) generally confers a high affinity and favourable selectivity towards αvβ6, with additional flanking residues further enhancing these properties^43,44^.

Amino acid modifications, peptide cyclization, and multimerization have also proven effective strategies to enhance peptide properties and functionality^45^. In particular, multimeric cyclic RGD (cRGD) peptide derivatives exhibited greater receptor affinity, increased tumor uptake, improved tumor-to-background (T/B) ratios and prolonged tumor retention compared to their monomeric counterparts^46, 47^. Furthermore, the avidity of these multivalent constructs has been shown to correlate directly with the number of incorporated cRGD units, resulting in a higher probability of receptor engagement and stronger overall binding. These findings were similarly observed during the development of the αvβ6-targeted trimers ^68^Ga-TRAP(SDM17)_3_^31^ and ^68^Ga-Trivehexin^48^, both based on the multifunctional and Gallium-68-selective chelator TRAP used as core scaffold. In particular, the promising preclinical performance of [^68^Ga]Ga-Trivehexin prompted early clinical investigations, which have convincingly demonstrated its utility for PET imaging of primary tumors and metastases with high target selectivity^49^. Its low intestinal uptake has proven beneficial in better delineation of metastases than alternative radiotracers based on alternative αvβ6-integrin binding peptides^26, 50^. To date, ^68^Ga-Trivehexin has been successfully applied in clinical PET/CT imaging across a broad spectrum of indications, including pancreatic ductal adenocarcinoma (PDAC)^51, 52^, thyroid cancer^53^, parathyroid adenoma^54^, pulmonary mucoepidermoid carcinoma^55^, head-and-neck squamous cell carcinoma (HNSCC)^56^ and its brain metastases^57^, non-small-cell lung cancer (NSCLC)^58^, as well as lobular and lymphatically metastasized breast cancer^59^. These studies collectively demonstrate the potential of ^68^Ga-Trivehexin as a versatile imaging agent for αvβ6-expressing malignancies.

Despite the undeniable utility of ^68^Ga-Trivehexin for these clinical indications, the relatively rapid radioactive decay of Gallium-68 (t_1/2_: 68 min) limits the applicable PET imaging window to approximately 2 to 3 hours p.i. (2 to 3 half-lives). In contrast, Zirconium-89 is a radiometal with a considerably longer half-life (t_1/2_: 78.4 h), thereby allowing the evaluation of pharmacokinetic properties and PET imaging potential at later time points^60, 61^. However, the TRAP chelator used for ^68^Ga-Trivehexin cannot be labelled with Zirconium-89.

Fusarinine C (FSC) is a multifunctional chelator that, like TRAP, provides a C3-symmetrical trimeric scaffold amenable to analogous functionalization strategies and can be efficiently labelled with Gallium-68. Notably, FSC additionally enables radiolabelling with Zirconium-89, thus providing a versatile platform for both short- and long-lived PET radionuclides.

FSC has previously been employed to develop RGD-based trimers targeting αvβ3 integrin^62–64^, and comparative studies with TRAP-based analogues revealed almost no significant differences in *in vivo* performance^65^.

In this study, we aimed to investigate whether a FSC-based αvβ6-targeted trimer could serve as a viable alternative to Trivehexin. To this end, we synthesized, radiolabeled, and characterized the ligand with both Gallium-68 and Zirconium-89, the latter being used for the first time to image αvβ6-expressing carcinomas. Its *in vitro* binding affinity, internalization ability, and *in vivo* imaging properties in αvβ6-expressing tumor models were systematically assessed. A direct comparison with ^68^Ga-Trivehexin was performed to evaluate whether the FSC scaffold preserves favorable pharmacokinetics and imaging performance, while providing the added advantage of late-time point imaging enabled by Zirconium-89.

## RESULTS

### Synthesis of the labelling precursor

The siderophore Fusarinine C (FSC) was extracted from *Aspergillus fumigatus* ΔsidG cultures following a previously published protocol^66^ and complexed with iron to prevent its hydroxamate groups from undergoing side reactions during the synthesis. Detailed procedures for the synthesis of the labelling precursor are provided in the Supplementary Information (**Figure S1-10**). Derivatisation of the three amino groups of [Fe]FSC with Azido-PEG4-Acid linkers was carried out via classic amide coupling with good yield (56.9%). Subsequently, three units of RGD targeting peptides were introduced by Cu(I)-catalyzed azide–alkyne cycloaddition (CuAAC) click reaction. Eventually, the coordinated metal was removed by transchelation leading to the labelling precursor with low yield (13.4%) (**Figure 1A**). The molecular design adopted was inspired by that used for Trivehexin (**Figure 1B**). Structurally, the two compounds differ for the chelator employed, and for the linkers considered for its functionalization. For the FSC-based trimer, PEG4 linkers were introduced to enhance the hydrophilicity of the final compound and to compensate for the higher lipophilicity of the chelator Fusarinine C compared to TRAP.

**Fig.1.**
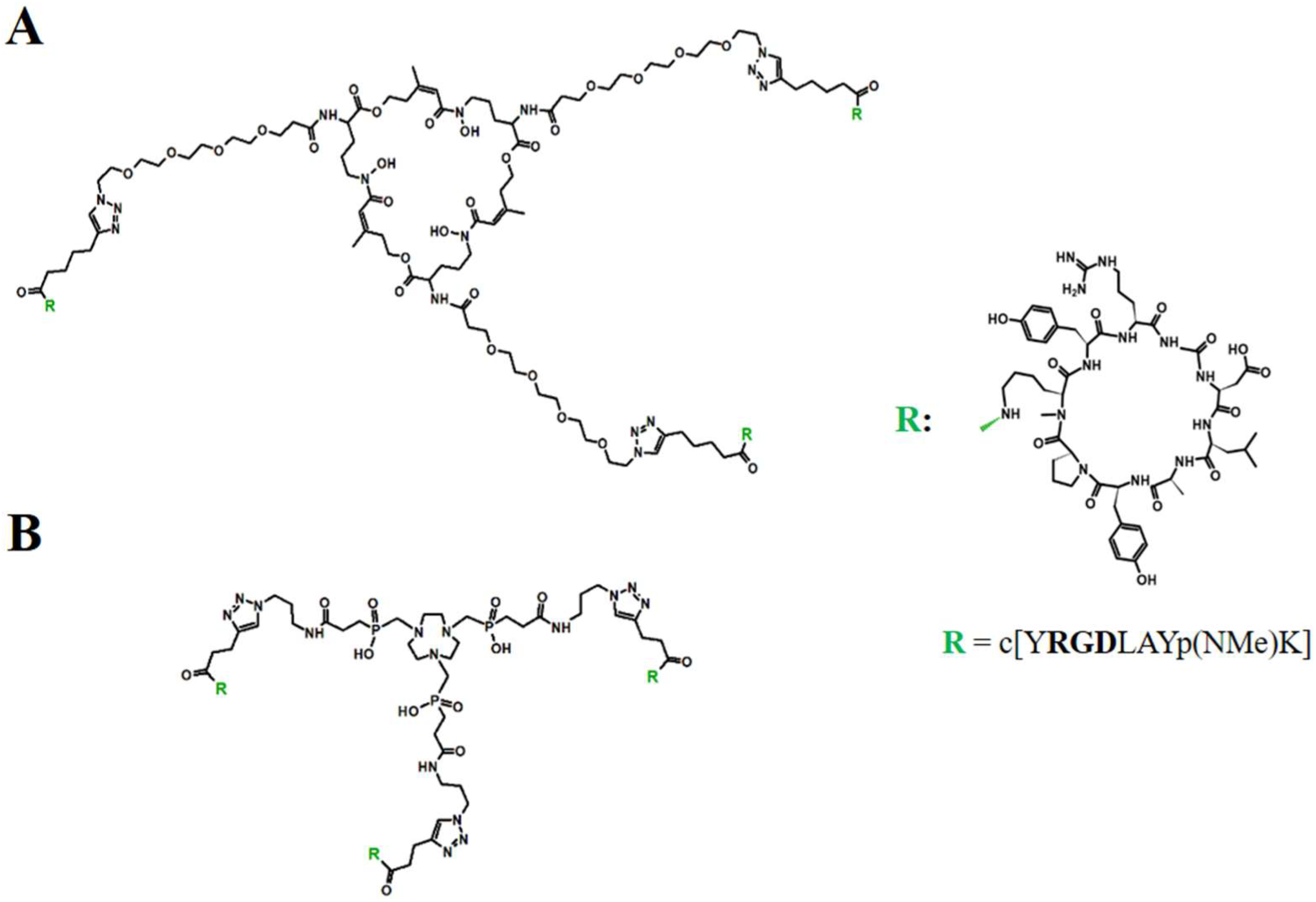
Chemical structures of the conjugates: (**A**) structure of the trimer FSC(PEG4-αvβ6)_3_ (MW: 5105.7 g/mol), (**B**) structure of Trivehexin (MW: 4300.6 g/mol).

### Radiolabelling with Gallium-68/Zirconium-89

The precursor FSC(PEG4-αvβ6)_3_ could be quantitatively labelled with Gallium-68 achieving a radiochemical yield (RCY) of > 98% as determined by radio-iTLC, and a radiochemical purity (RCP) of > 97%. Importantly, the labelled compound could be used directly in all experiments without further purification (**Figure S11A-12A**). A molar activity of up to 99.5 MBq/nmol was obtained. Notably, the radiolabelling of this precursor could be completed within 10 minutes at room temperature, demonstrating high labelling efficiency at the mild conditions applied.

We have also been able to achieve a similarly high RCY of 99.2% (determined by radio-iTLC) and a RCP of > 98.5% for Zirconium-89 labelling, performed at 40°C (**Figure S11B-12C**). However, in this case, the maximum molar activity obtained was limited to 2.0 MBq/nmol (without optimization; **Figure S13**).

### *In vitro* characterization

First, we determined the affinity of FSC(PEG4-αvβ6)_3_ and of [^nat^Ga]Ga-FSC(PEG4-αvβ6)_3_ for the αvβ6 integrin, using a well-established ELISA assay ^67, 68^. For the unlabelled compound, we obtained an IC*_50_* value of 0.69 ± 0.24 nM, confirming the subnanomolar affinity observed for other Tyr2-based multimers^69^. The ^nat^Gallium-labelled version showed an approximately two-fold higher IC*_50_* of 1.53 ± 0.62 nM, indicating a lower affinity compared to [^nat^Ga]Ga-Trivehexin (0.048 nM)^48^ (**Figure S14**, **Table 1A**).

**Table 1.**
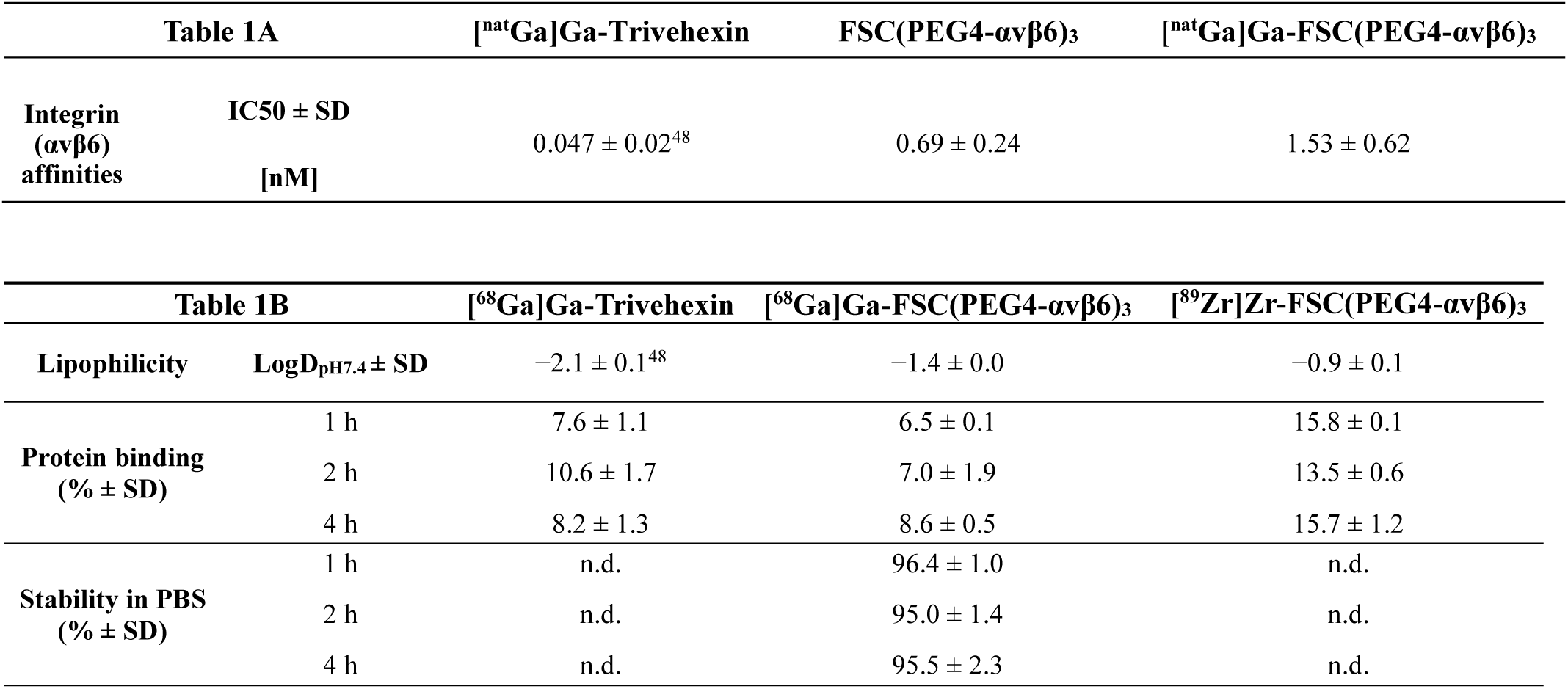
(**A**) αvβ6-binding affinity of FSC(PEG4-αvβ6)_3_ and [^nat^Ga]Ga-FSC(PEG4-αvβ6)_3,_ assessed by determining the half-maximal inhibitory concentration (IC50). The IC50 value for [^nat^Ga]Ga-Trivehexin was obtained from literature^48^. (**B**) Results of lipophilicity (LogD_pH7.4_), protein binding as well as stability determination in human serum (% of intact radiotracer) for Gallium-68 labelled FSC(PEG4-αvβ6)_3_, Trivehexin and for [^89^Zr]Zr-FSC(PEG4-αvβ6)_3_.

[^68^Ga]Ga-FSC(PEG4-αvβ6)_3_ showed lower hydrophilicity when directly compared to [^68^Ga]Ga-Trivehexin (Log D_7.4_= -1.4 ± 0.1 vs -2.1 ± 0.1^48^) (**Figure S15**, **Table 1B**). This could be due to the influence of the larger chelator Fusarinine C, which was only partially compensated by the PEG4 linkers. The distribution coefficient found for [^89^Zr]Zr-FSC(PEG4-αvβ6)_3_ was moderately lower (Log D_7.4_= -0.9 ± 0.1) than that of the Gallium-68 version. This finding was contrary to the expectations, given that the Zirconium (Zr^4+^) complex is mono-charged, whereas the Gallium (Ga^+3^) version is neutral. A rationale for this result remains to be elucidated.

For both FSC-based radiocompounds, the affinity to serum proteins was overall low and remained consistent over time (**Figure S16**, **Table 1B**). An approximately 2-fold higher serum protein binding was observed for [^89^Zr]Zr-FSC(PEG4-αvβ6)_3,_ compared to [^68^Ga]Ga-FSC(PEG4-αvβ6)_3_.

The stability of [^68^Ga]Ga-FSC(PEG4-αvβ6)_3_ in PBS and of [^89^Zr]Zr-FSC(PEG4-αvβ6)_3_ in the labelling solution was determined over time, and no significant release of the radionuclide or degradation of the radiocompound was observed (**Figure S17A**, **Table 1B**). Cell binding studies were conducted for [^68^Ga]Ga-FSC(PEG4-αvβ6)_3_ in αvβ6-expressing H2009 cells as well as in αvβ6-negative MDA-MB-231 cells (**Figure 2A**). The results confirmed highly specific αvβ6-mediated uptake, which was efficiently inhibited by more than 89% upon co-incubation with a 1000-fold excess of the c[YRGDLAYp(NMe)K]-alkyne peptide in H2009 cells.

**Fig.2.**
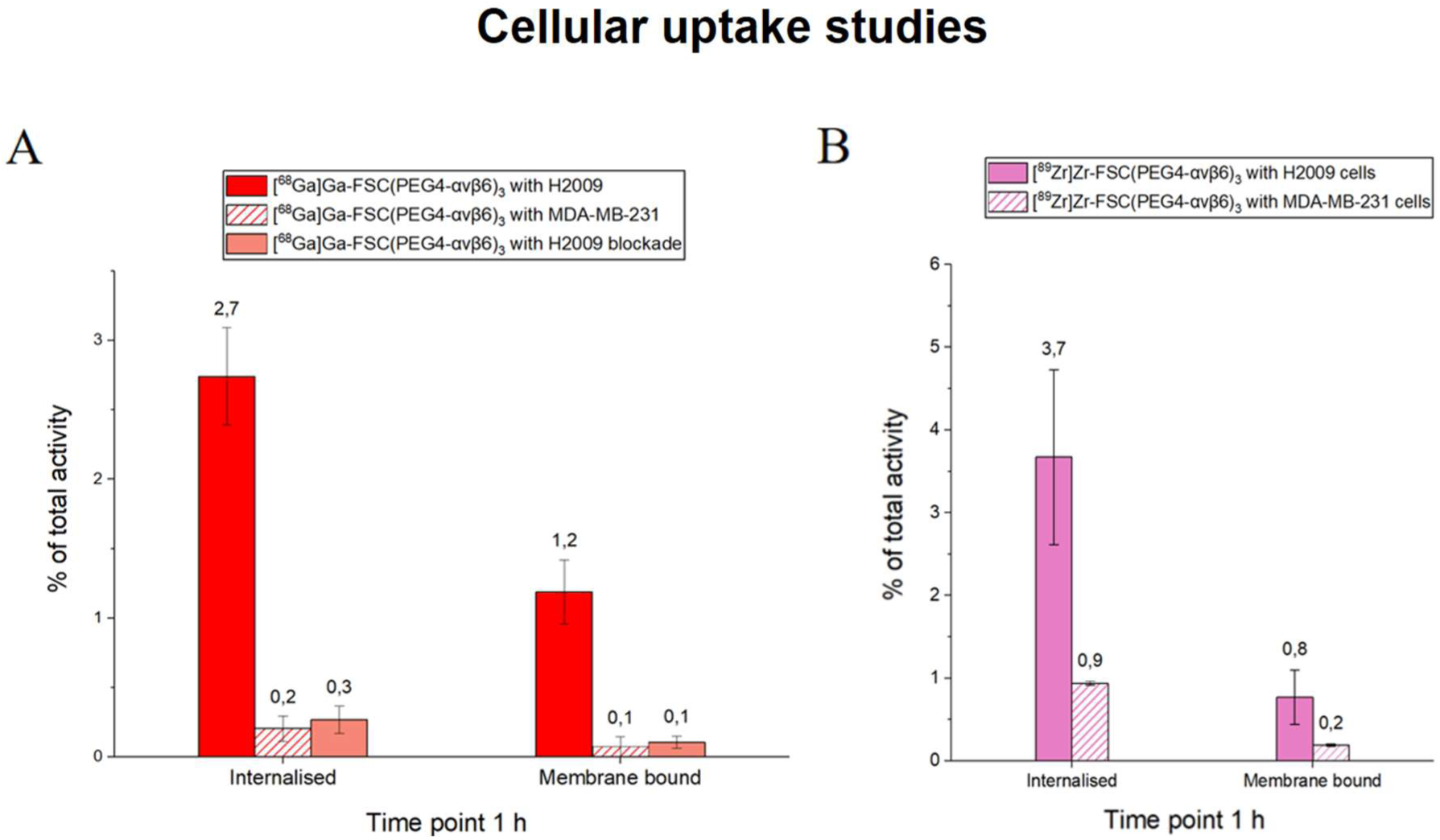
Cellular uptake studies using H2009 (αvβ6 positive human lung adenocarcinoma) and MDA-MB-231 (αvβ6 negative human breast cancer) cell lines. (**A**) Cellular uptake for [^68^Ga]Ga-FSC(PEG4-αvβ6)_3_. Blocking was performed with a 1000-fold excess of c[YRGDLAYp(NMe)K]-alkyne peptide. The results are presented as mean values of three independent experiments. (**B**) Cellular uptake for [^89^Zr]Zr-FSC(PEG4-αvβ6)_3_. The results are presented as mean values derived from two independent experiments.

In contrast, uptake in αvβ6-negative MDA-MB-231 cells was negligible, accounting for less than 0.3% of the total activity, thereby highlighting the high selectivity for αvβ6-positive cells. Specific receptor-mediated internalization was also obtained for the compound [^89^Zr]Zr-FSC(PEG4-αvβ6)_3_ with higher nonspecific binding values, but overall, no significant difference to the Gallium-68 counterpart (**Figure 2B**) was observed. In this case, nonspecific internalization was assessed exclusively in αvβ6-negative MDA-MB-231 cells, without using blocking conditions.

### *In vivo* characterization

The biodistribution profile of [^68^Ga]Ga-FSC(PEG4-αvβ6)_3_ was preliminarily evaluated in healthy BALB/c mice (**Figure S18**). The results showed fast clearance from the blood pool and negligible nonspecific accumulation. The high renal activity known from [^68^Ga]Ga-Trivehexin^48^ (70.6 ± 20.0 % ID/g at 90 min p.i. in MDA-MB-231 bearing SCID mice) was also present in [^68^Ga]Ga-FSC(PEG4-αvβ6)_3_ (72.6 ± 3.2 % ID/g at 90 min p.i.). Investigation of the *in vivo* metabolism of [^68^Ga]Ga-FSC(PEG4-αvβ6)_3_ at a time point of 15 minutes p.i. revealed high stability in murine serum, whereas evidence of degradation and partly release of Gallium-68 was observed in urine (**Figure S19**).

In H2009 tumor-bearing mice, the head-to-head comparison via PET imaging at a time point of 75-90 min p.i. showed clear tumor delineation with both, [^68^Ga]Ga-Trivehexin (2.3 ± 0.2% ID/g PET quantification) and [^68^Ga]Ga-FSC(PEG4-αvβ6)_3_ (1.4 ± 0.2 % ID/g), and low uptake in other organs with the exception of the kidney (**Figure 3A-B**). Uptake specificity could be confirmed in the blocking group, where tumor uptake of [^68^Ga]Ga-FSC(PEG4-αvβ6)_3_ was not discernible (**Figure 3A**). Kidney uptake of both radiotracers was found in a similar range (22-35 % ID/g) via PET quantification (**Figure 3C-D**).

**Fig.3.**
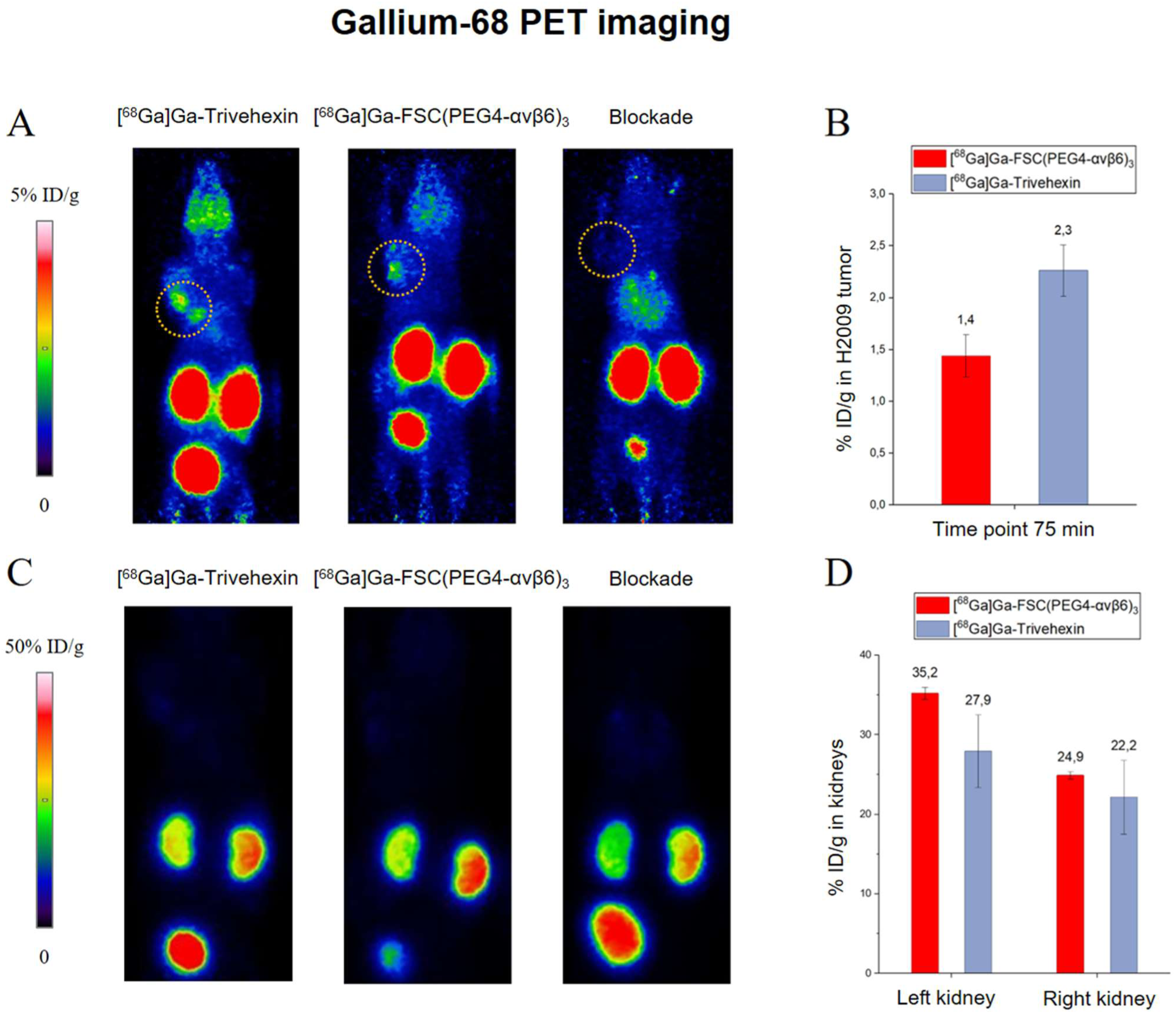
Gallium-68 PET imaging results using H2009-bearing SCID mice. (**A**) Maximum intensity projection (MIP) PET images at 75-90 min p.i. of [^68^Ga]Ga-Trivehexin (following an injected amount of 68 pmol, 4.6 MBq), [^68^Ga]Ga-FSC(PEG4-αvβ6)_3_ (amount injected 139 pmol, 7.2 MBq) and [^68^Ga]Ga-FSC(PEG4-αvβ6)_3_ (amount injected 74 pmol, 3.3 MBq) under blocking conditions. The tumor position is indicated with a dashed circle. The same xenografted animal was used in the case of the scan following injection of [^68^Ga]Ga-FSC(PEG4-αvβ6)_3_ and [^68^Ga]Ga-Trivehexin, with a 3 days recovery period between the injections. (**B**) Comparison of [^68^Ga]Ga-FSC(PEG4-αvβ6)_3_ or [^68^Ga]Ga-Trivehexin uptake in H2009 tumors bearing animals. Values were calculated in manually drawn ROIs, fitted to the tumors. Results are expressed as mean % ID/g values (n=3). (**C**) MIP PET images (same as in A) at 75–90 min p.i., scaled to kidney uptake. (**D**) Comparison of uptake in left and right kidney of animals injected with [^68^Ga]Ga-FSC(PEG4-αvβ6)_3_ or [^68^Ga]Ga-Trivehexin. Values were calculated in manually drawn ROIs fitting to each kidney. Results are expressed as mean % ID/g values (n=3).

The biodistribution data showed the typical biodistribution profile of c[YRGDLAYp(NMe)K]-based radiopharmaceuticals with fast blood clearance, and blockable uptake in the αvβ6-expressing tissues (stomach and intestines) in addition to the xenograft tumor, where blocking reduced the tumor uptake from 1.4 ± 0.4 % ID/g to 0.4 ± 0.1% ID/g (**Figure 4A**). Blocking also slightly reduced kidney uptake from 73.7 ± 4.7 % ID/g to 51.1 ± 2.5 % ID/g. As an unexpected finding, lung uptake of [^68^Ga]Ga-FSC(PEG4-αvβ6)_3_ was fourfold increased in the blocking group, which has not been observed before with other radiopharmaceuticals based on the same peptide sequence. The underlying reason for this discrepancy remains unclear. The target-to-organ ratios observed in our study were consistently high, exceeding a factor of 5 for blood, pancreas, heart, adrenals, and muscle. These values indicate a highly favorable biodistribution profile, confirming a rapid clearance from the circulation and minimal nonspecific uptake in critical organs (**Figure 4B**).

**Fig.4.**
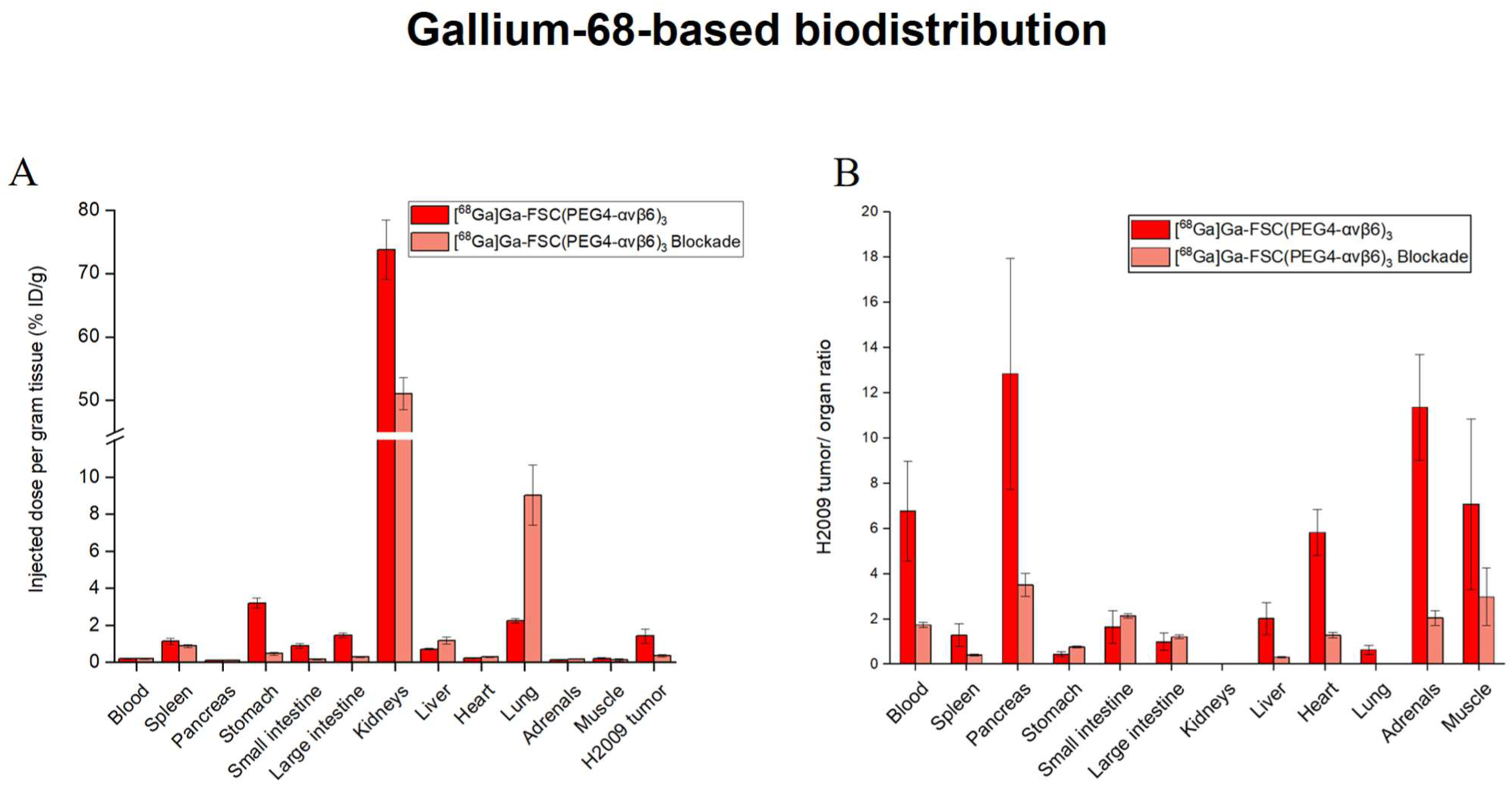
Gallium-68-based biodistribution studies. (**A**) Ex vivo biodistribution experiments in H2009-bearing SCID mice (n = 3), performed 90 min p.i. for [^68^Ga]Ga-FSC(PEG4-αvβ6)_3_ (amount injected 120-140 pmol, 6.8-8.0 MBq), and for [^68^Ga]Ga-FSC(PEG4-αvβ6)_3_ (amount injected 73-80 pmol, 5.2-5.7 MBq) under blocking conditions (pre-injection of 50 nmol of unlabelled Trivehexin 10 min prior to radiotracer). (**B**) Tumor-to-organ ratios derived from H2009 biodistribution data for [^68^Ga]Ga-FSC(PEG4-αvβ6)_3_.

[^89^Zr]Zr-FSC(PEG4-αvβ6)_3_ was also able to delineate the H2009 xenografts via PET imaging (**Figure 5**). Tumor uptake was clearly visible at time points of 75 min, 1 day and 3 days p.i. and to a lesser extent at 6 days p.i. with a low background in other organs and the typical high kidney uptake. Starting at the 1-day time point, bone uptake was visible in PET scans, suggesting initial degradation of the radioligand and subsequent release of free Zirconium-89, which accumulated in the joints. Image quality of the Zirconium-89 scans was negatively affected by the low injected activity of 0.2-0.4 MBq per mouse, which was necessary to avoid target saturation and blocking as a result of the relatively low molar activity achieved (2 MBq/nmol, **Figure S13**). Comparison of the two FSC-based ligands revealed that [^89^Zr]Zr-FSC(PEG4-αvβ6)_3_ achieved tumor uptake comparable to [^68^Ga]Ga-FSC(PEG4-αvβ6)_3_ at 90 min p.i. (% ID/g:1.1 ± 0.2 vs 1.4 ± 0.4, respectively, **Figure 4A** and **6A**). Uptake in non-target organs was likewise similar, indicating that the longer-lived Zirconium-89 label does not substantially alter the biodistribution profile at early time points. However, blood clearance of [^89^Zr]Zr-FSC(PEG4-αvβ6)₃ was slower, compared to its Gallium-68 analogue, as reflected by significantly higher blood activity at 90 min post-injection (%ID/g: 0.8 ± 0.1 vs. 0.2 ± 0.0, respectively). Notably, T/O ratios for the Zirconium-89–labelled compound steadily increased over time in most organs, including blood, intestine, heart and muscle, up to 6 days post-injection, indicating a prolonged tumor retention relative to normal tissues (**Figure 6B**). In contrast, no such trend was observed in the stomach and liver, where T/O ratios remained relatively stable.

**Fig.5.**
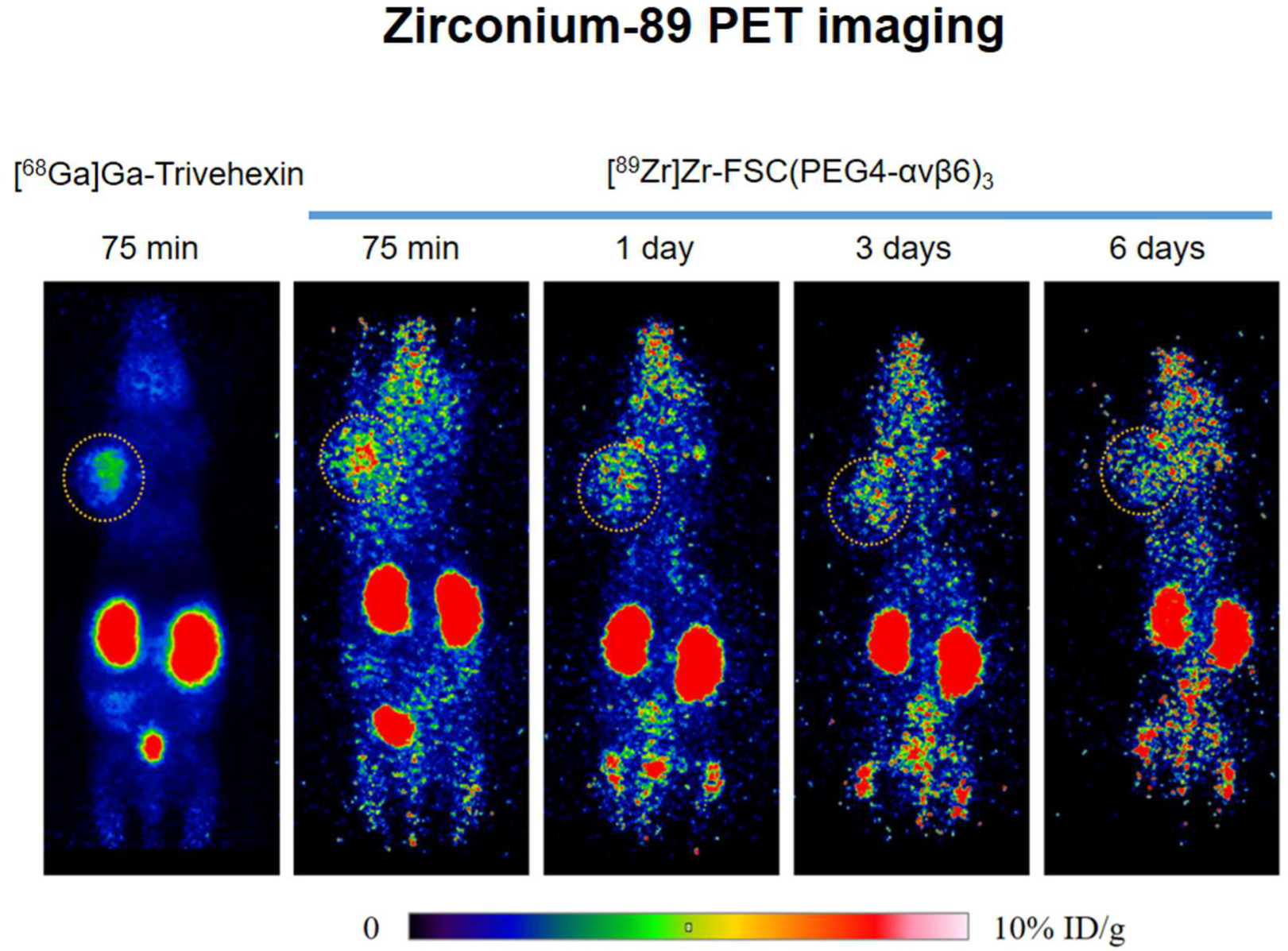
MIP PET image of [^68^Ga]Ga-Trivehexin at a time point of 75 min p.i. (amount injected, 72 pmol, 3.8 MBq) and of [^89^Zr]Zr-FSC(PEG4-αvβ6)_3,_ at different time points from 75 min up to 6 days p.i. (200 pmol, 0.4 MBq), obtained using the same xenografted animal, with 1 day recovery period. The tumor position is indicated with a dashed circle.

**Fig.6.**
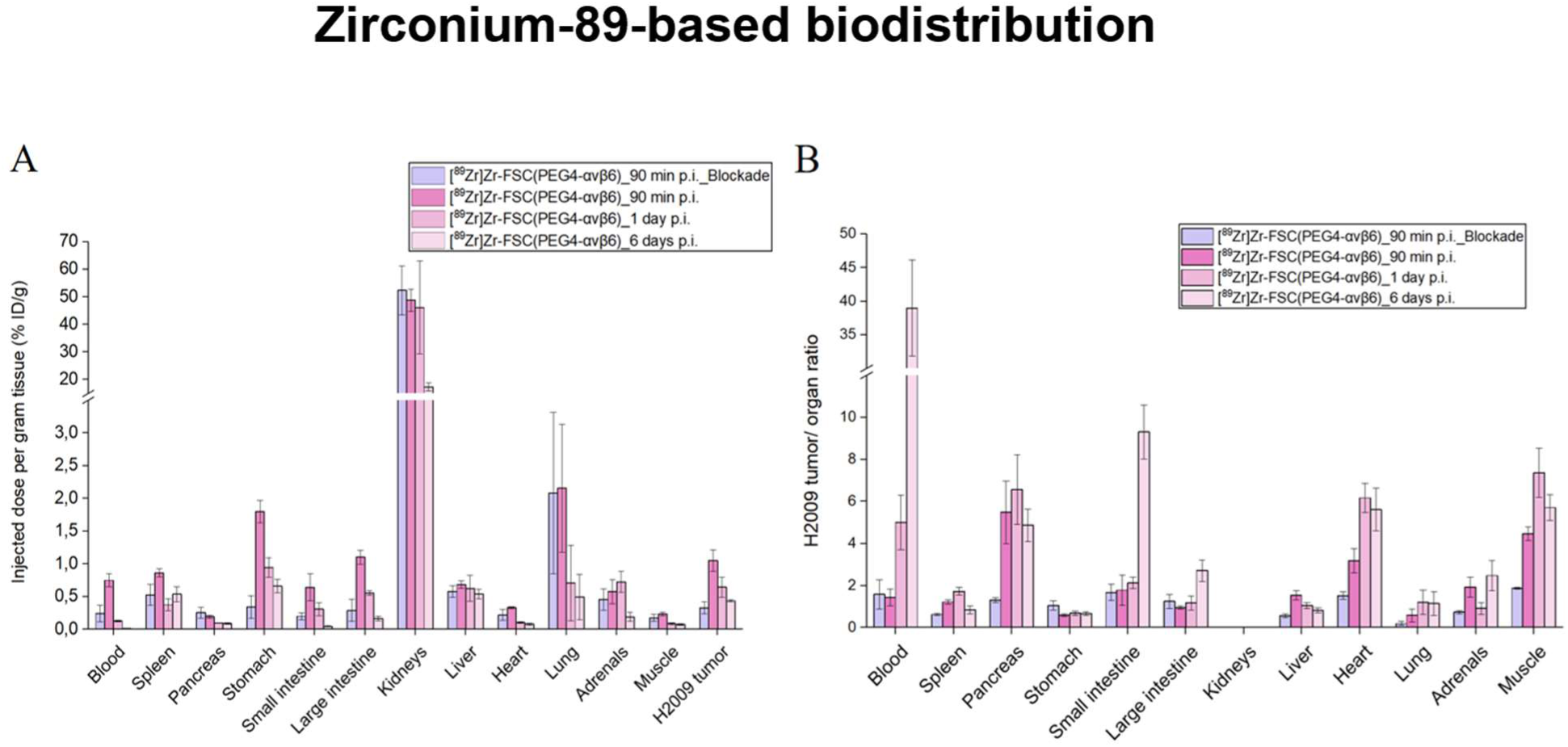
Zirconium-89-based biodistribution studies. (**A**) Ex vivo biodistribution experiments in H2009-bearing SCID animals (n = 3) performed at different time points from 90 min up to 6 days p.i. for [^89^Zr]Zr-FSC(PEG4-αvβ6)_3_ (amount injected 200-400 pmol, 0.4-0.8 MBq) and for [^89^Zr]Zr-FSC(PEG4-αvβ6)_3_ (amount injected, 260-320 pmol, 0.5-0.6 MBq) under blocking conditions (pre-injection of 50 nmol of unlabelled Trivehexin 10 min prior to radiotracer). (**B**) Tumor-to-organ ratios derived from H2009 biodistribution data for [^89^Zr]Zr-FSC(PEG4-αvβ6)_3_.

Hence, the newly developed FSC-based trimer FSC(PEG4-αvβ6)_3_ accumulates with high specificity in αvβ6-expressing tumors and is suitable to investigate ligand pharmacokinetics at delayed time points using Zirconium-89.

## DISCUSSION

The cyclic nonapeptide c[YRGDLAYp(NMe)K] represents the result of extensive optimization efforts aimed at downsizing and metabolically stabilizing the RGD motif derived from the foot-and-mouth disease virus (FMDV) peptide^45^. This sequence has been identified as a stable, high-affinity, and αvβ6-selective ligand. Furthermore, its terminal lysine side-chain group is amenable to conjugation, making this peptide an attractive platform for the development of clinically relevant targeted tracers. To further improve *in vivo* target interaction, multimerization strategies were explored, using the TRAP chelator as a scaffold. This optimization lead to the synthesis of [^68^Ga]Ga-Trivehexin^48^. Compared to an analogue trimer candidate in which both tyrosines were substituted with phenylalanines, [^68^Ga]Ga-Trivehexin exhibited a moderate increase in hydrophilicity but drastically improved pharmacokinetic properties, including rapid blood clearance and reduced nonspecific uptake in most organs, with the kidneys representing the sole exception^48^.

Reducing this kidney uptake could be favorable for clinical translation and adoption. In a study from our group^70^, we directly compared two analogous dual-modality imaging agents targeting the Cholecystokinin-2 receptor (CCK2R), differing only in the chelator scaffold (TRAP vs. Fusarinine C). In this study, we demonstrated that the chelator had largely comparable influence on *in vitro* and *in vivo* performance, with the exception of higher renal accumulation and retention observed for the TRAP-based construct, which was significanty greater by approximately 60% at 120 min p.i. In other previous works, we have also demonstrated the utility of Fusarinine C as a versatile platform for the development of multimers targeting the integrin αvβ3^62–64^. Building on these experiences and motivated by the promising preclinical and clinical performance of [^68^Ga]Ga-Trivehexin, we sought to develop and investigate an analogous tracer that maintains the peptide sequence, valency, and geometry of Trivehexin while employing Fusarinine C as scaffold. This design, while enabling the additional labelling with the longer-lived radionuclide Zirconium-89, aimed to assess whether such a probe could provide comparable or even improved targeting and pharmacokinetic properties, particularly lower renal accumulation, thereby supporting its potential as a platform for the future development of αvβ6-targeted radiopharmaceuticals.

Modifications to the polarity of the peptide sequence have led to dramatic and unexpected improvements in pharmacokinetic properties, in particular in the development of Trivehexin. Therefore, in designing a new candidate, we carefully considered the overall molecular polarity and decided to introduce 3 flexible PEGylated linkers, which aimed at increasing the hydrodynamic radius and thus reducing nonspecific binding^71^. In addition, PEGylation can extend the spatial distance between individual binding units, which may increase the avidity of multimeric ligands. This could facilitate simultaneous binding to multiple targets and hence improve the tumor retention of the tracer *in vivo*^72^. As a result, FSC(PEG4-αvβ6)₃ exhibits an approximately 19% increase in molecular weight, compared to Trivehexin, a change we deemed acceptable given the expected benefits in hydrophilicity and pharmacokinetic behaviour.

Despite this structural modification, our data indicated that [^68^Ga]Ga-FSC(PEG₄-αvβ6)₃ (LogD_pH7.4_ = −1.4 ± 0.0; Rt: 13.5 min) is less hydrophilic than [^68^Ga]Ga-Trivehexin (LogD_pH7.4_ = −2.1 ± 0.1; Rt: 12.8 min; **Table 1B**, **Figure S11A** and **Figure S15**). The results indicated that the PEGylated spacers did not compensate for the lower polarity of the Fusarinine C chelator, compared to TRAP.

Still, [^68^Ga]Ga-FSC(PEG₄-αvβ6)₃ exhibited similar low affinity for human serum proteins, compared to [^68^Ga]Ga-Trivehexin (**Table 1B**). Notably, comparison of the *in vivo* biodistribution profiles revealed that [^68^Ga]Ga-FSC(PEG₄-αvβ6)₃ exhibited more rapid blood clearance and at least 50% lower accumulation in most non-target organs, with the exception of liver, stomach and kidneys (**Figure S20**). The introduction of PEGylated linkers therefore improved pharmacokinetics by reducing nonspecific binding in organs such as the lungs and intestines, which are frequent sites of primary lesions and metastases in αvβ6-integrin–expressing carcinomas.

Regarding the interaction of [^68^Ga]Ga-FSC(PEG4-αvβ6)₃ with its target, cellular uptake studies showed slightly inferior binding of the new candidate to αvβ6-expressing cells, compared to the results reported for [^68^Ga]Ga-Trivehexin (% of H2009 cell associated activity: 3.9 vs 4.9, respectively **Figure 2A**)^48^. In line with this, the affinity of [^nat^Ga]Ga-FSC(PEG₄-αvβ6)₃ was also lower (though still in the nanomolar range), compared to [^nat^Ga]Ga-Trivehexin, which exhibited an IC50 in the double-digit picomolar range^48^ **(Table 1A and Figure S14)**. Accumulation in H2009 xenografts for [^68^Ga]Ga-FSC(PEG4-αvβ6)₃ was limited to 1.4 % ID/g, as determined by both, *ex vivo* biodistribution (**Figure 4A**) and PET imaging quantification (**Figure 3B**). This value was 1.6-fold lower than observed for [^68^Ga]Ga-Trivehexin (2.3 % ID/g) in quantitative PET analysis performed in the same group of mice. This could be explained by the lower target affinity of [^68^Ga]Ga-FSC(PEG₄-αvβ6)₃ and may also reflect the effect of the increased distance between the individual peptidic units. This is consistent with previous findings reporting a moderate reduction in affinity associated with the introduction of PEG10 spacer in αvβ6-targeted TRAP trimers^73^. These results underscore the need for further studies to elucidate how PEGylated linkers of varying lengths influence both, specific target binding and nonspecific interactions, as these effects may be governed by different mechanisms.

Notably, the level of tumor accumulation for [^68^Ga]Ga-Trivehexin was considerably lower than in previous studies, and approximately 3.3-fold lower than the value reported in the literature for biodistribution studies^48^ (**Figure S20**). This discrepancy may reflect lower expression of αvβ6-integrin in the H2009 xenografted tumors of our cohort, however we did not investigate this aspect further. Overall, even though [^68^Ga]Ga-FSC(PEG4-αvβ6)₃ showed impaired tumor targeting as compared to [^68^Ga]Ga-Trivehexin, this was compensated by the lower off target accumulation particular in relevant organs and blood.

Beyond its well-established applications in immuno-PET, Zirconium-89 presents a compelling opportunity for labelling proteins and peptides with prolonged *in vivo* circulation, such as αvβ6-targeted multimers, which demonstrated tumor retention over several days^69^. In such situations, the long physical half-life of Zirconium-89 allows for longitudinal imaging, which is particularly valuable in preclinical studies to obtain detailed *in vivo* data on biodistribution and tissue clearance at late time points. From a clinical perspective, delayed imaging with Zirconium-89 provides a superior ability to detect lesions with low ligand avidity, which are often difficult to visualize using tracers labeled with short-lived radionuclides. Indeed, first-in-human studies with [⁸⁹Zr]Zr-DFO-PSMA demonstrated the localization of 15 PSMA-positive lesions across eight patients, with a significantly higher SUVmax compared to initial PET scans^74^. These results underscore the potential of [⁸⁹Zr]Zr-labelled tracers not only for improved lesion detection, but also for providing more accurate assessments of target engagement, supporting their broader application in both drug development and patient stratification.

Previous studies from our group demonstrate the feasibility of using Fusarinine C as a cyclic chelator for Zirconium-89 labeling, as well as its superior stability and kinetic inertness, compared to DFO^61, 75^. These preliminary studies demonstrate that the Zirconium-89 tracer exhibited biological properties comparable to its Gallium-68 counterpart, with even higher tumor-to-blood ratios. Based on these considerations, we also investigated the potential of [^89^Zr]Zr-FSC(PEG₄-αvβ6)₃ as a tracer to evaluate pharmacokinetics at later time points.

In contrast to previous results and to our expectations based on the overall +1 charge of the [^89^Zr]Zr-FSC complex, compared to the neutral charge of the [^68^Ga]Ga-FSC complex, [^89^Zr]Zr-FSC(PEG4-αvβ6)₃ exhibited a less negative LogD_pH7.4,_ compared to its Gallium-68 counterpart (LogD_pH7.4_ = −0.9 ± 0.1; Δ = 0.5; **Table 1B**, **Figure S15**), while showing comparable HPLC retention time (**Figure S11B**).

This presumed lower polarity of [⁸⁹Zr]Zr-FSC(PEG4-αvβ6)₃ was consistent with its approximately two-fold higher affinity for serum proteins, compared to [⁶⁸Ga]Ga-FSC(PEG4-αvβ6)₃ (**Table 1B**), as well as with the increased nonspecific binding to αvβ6-negative cells observed in cell uptake studies (**Figure 2B**). Despite an overall slower clearance—in line with the *in vitro* findings and reflected by higher blood persistence and lower renal levels—the *in vivo* biodistribution of [⁸⁹Zr]Zr-FSC(PEG4-αvβ6)₃ was largely comparable to that of [⁶⁸Ga]Ga-FSC(PEG4-αvβ6)₃ (**Figure 4A** and **Figure 6A**). Moreover, tumor uptake in H2009 xenografts was similar for both tracers. In contrast to previously reported tracers, which could be labelled up to 25 MBq/nmol^61^, [⁸⁹Zr]Zr-FSC(PEG4-αvβ6)₃ reached a maximum molar activity of only 2.0 MBq/nmol under non-optimized conditions (**Figure S13**). The relatively low molar activity achievable for [⁸⁹Zr]Zr-FSC(PEG4-αvβ6)₃, combined with the need to inject modest tracer amounts into the xenografted mice to avoid potential target saturation, resulted in low injected doses (0.4-0.8 MBq), which negatively affected overall image quality (**Figure 5**). Nevertheless, the results obtained from the preclinical imaging studies still allowed satisfactory tumor identification, and provided valuable pharmacokinetic information indicating that 41.6 % ID/g of the initial accumulation observed at 90 min p.i. was retained at 6 days p.i. (**Figure 6A**). The PET images revealed a noticeable increase in uptake in the joints and spine, starting at 1 day p.i., suggesting partial release of Zirconium-89 from the ligand. This behavior contrasts with previous imaging results of other [⁸⁹Zr]Zr-FSC-based tracers at 1 day p.i., which showed prolonged *in vivo* stability of the complex^61^. This limited *in vivo* stability may reflect the incomplete saturation of the Zirconium^4+^ coordination sphere-which requires eight donor groups-by the hexadentate Fusarinine C.

## CONCLUSIONS

In this study, we report the first development of an αvβ6-targeted trimeric tracer based on Fusarinine C and its evaluation with both Gallium-68 and Zirconium-89. The novel tracer demonstrated overall satisfactory interaction with αvβ6, albeit with lower affinity and cell internalization, compared to [^68^Ga]Ga-Trivehexin. Interestingly, this reduced target interaction was accompanied by lower nonspecific binding, an effect we attribute primarily to the introduction of PEGylated spacers. These findings highlight the critical role of spacer design in the optimization of αvβ6-targeted tracers, and suggest that further refinement of this element could enhance future derivatives. While some indications of *in vivo* degradation of [^89^Zr]Zr-FSC(PEG₄-αvβ6)₃ were observed, its performance still supported valuable preclinical assessment of pharmacokinetics and biodistribution at late time points, suggesting its potential for extended imaging studies. Overall, our results provide an important foundation for the continued development of next-generation αvβ6-targeted diagnostic agents.

## EXPERIMENTAL SECTION

### Instrumentation

#### Analytical [radio]-RP-HPLC

RP-HPLC analysis was performed on a UltiMate 3000 system equipped with pump, autosampler, column compartment, diode array detector (Thermo Fisher Scientific, Vienna, Austria) and radio detector (GabiStar, Raytest; Straubenhardt, Germany).

##### Method

A Jupiter 4 μm Proteo 90 Å 250 x 4.6 mm (Phenomenex Ltd. Aschaffenburg, Germany) column with a flow rate of 1 mL/min and UV detection at 220 nm was used. Acetonitrile (ACN)/H_2_O + 0.1% trifluoroacetic acid (TFA) was used as mobile phase with the following multistep gradient: 0.0-3.0 min 10% ACN, 3.0-16.0 min 10-60% ACN, 16.0-18.0 min 60% ACN, 18.0-18.1 min 60-10% ACN, 18.1-22.0 min 10% ACN.

#### ^68^[Ge]Ge/^68^[Ga]Ga-Generator

[^68^Ga]GaCl3 was obtained from a commercial 68Ge/68Ga generator (Eckert and Ziegler, Berlin, Germany) eluted with 0.1 N HCl solution (Rotem Industries, Dimona, Israel). The fractionated elution method was used in order to increase the ratio of activity to volume to its maximum (150-200 MBq in 1.5 mL).

#### γ-Counter

The 2480 Automatic Gamma counter Wizard2 3” (PerkinElmer Life Sciences and Analytical Instruments, formerly Wallac Oy, Turku, Finland) was used to measure the radioactivity of the samples.

#### Radio-iTLC

Radio instant thin layer chromatography (*radio*-ITLC) analysis of the Gallium-68 compounds were performed using iTLC-SG stripes (Agilent Technologies, Folsom, CA, USA) and 0.1 M sodium citrate solution (pH 5). The strips spotted with samples were analyzed using a TLC scanner (Scan-RAM, LabLogistic, Sheffield, UK). Gallium-68 labelled bioconjugates remained at the origin (Rf < 0.1), while the unbound radionuclide migrated to the solvent front (Rf > 0.9).

For Zirconium-89 labelling, iTLC-SG stripes were eluted with 0.05 M EDTA solution (pH7) and then analysed with with Cyclone Plus (Perkin Elmer, Waltham, US).

## MATERIALS AND METHODS

All commercially available chemicals, reagents and solvents were of analytical grade and were used without further purification. Only high-purity water (18 mΩ) was employed. Trivehexin was kindly provided by TRIMT GmbH. H2009 (CRL-5911) and MDA-MB-231 (HTB-26) were obtained from the American Type Culture Collection (ATCC), Manassas, VA, USA). 1M Zirconium-89 oxalic acid solution (1 MBq/µL) was purchased from Perkin Elmer (Waltham, US).

All other reagents were purchased from Sigma-Aldrich (Merck, KGaA, Darmstadt, Germany) or Merck (Darmstadt, Germany).

### Radiochemistry

For Gallium-68 labelling, 6 nmol of FSC(PEG4-αvβ6)_3_ were incubated with 200 µL eluate (30-40 MBq) and with 42 µL of 1.1 M sodium acetate solution (pH 8.8) to reach a final pH of 4.4. For Trivehexin, 3 nmol of precursor were mixed with 200 µL eluate and 20 µL of sodium acetate solution to reach a pH of 3. The labelling was performed within 10 min at RT and 95°C for the experimental FSC-based compound and Trivehexin respectively.

For Zirconium-89 labelling, 7 uL of Zirconium oxalic solution (7 MBq) were neutralized with 6.7 µL of 1 M NaCO_3_. After 3 min, 100 uL of 0.5 M HEPES buffer (pH 7) were added together with 7.5 nmol of FSC(PEG4-αvβ6)_3_ precursor. The mixture was incubated for 30 min at 40°C under shaking.

The purity of the radiolabelled compounds was determined both, by *radio*-RP-HPLC and by *radio*-iTLC. For *in vivo* experiments, [^68^Ga]Ga-labelling was carried out on a fully automated synthesis module (GallElut^+^, Scintomics GmbH, Gräfelfing, Germany) according to a previously described protocol [1]. 2 nmol Trivehexin or FSC(PEG4-αvβ6)_3_ were labeled with 300-500 MBq of Gallium-68 at pH=2.0.

The purity of the radiolabeled compounds was confirmed by *radio*-iTLC and *radio*-HPLC. For *radio*-iTLC, we used silica impregnated glass fiber chromatography paper (ITLC® by Agilent) as stationary phase and 0.1 M aq. sodium citrate as mobile phase (purities were > 95%). *Radio*-HPLC was performed on a Shimadzu RP-HPLC system including a NaI(Tl) well-type scintillation counter from Elysia-Raytest (type Gabi; Straubenhardt, Germany). A linear 10-90% gradient (MeCN/H_2_O both supplemented with 0.1% TFA) in 15 minutes was used.

Injected molar activities were 55-79 MBq/nmol for ^68^Ga-TVH and 29-89 MBq/nmol for ^68^Ga-FSC(PEG4-αvβ6)_3_, respectively.

Zirconium-89 labelling of FSC(PEG4-αvβ6)_3_ for *in vivo* experiments was conducted using 23.5 MBq of ^89^Zr^4+^ (oxalic acid solution), which was neutralized with 23 µl of Na_2_CO_3_ (1M) for 3 minutes, followed by addition of 150 µl (4-(2-hydroxyethyl)-1-piperazineethanesulfonic acid buffer (HEPES, 0.5 M, pH 7) and 11.75 nmol FSC(PEG4-αvβ6)_3_. The mixture was heated to 80°C for 30 minutes. The purity of [^89^Zr]Zr-FSC(PEG4-αvβ6)_3_ was confirmed by *radio*-TLC using ethylenediaminetetraacetic acid (EDTA, 0.05 M) as the mobile phase (purities were > 95%). The injected molar activity was 2 MBq/nmol.

### Distribution coefficient (LogD_pH7.4_), stability in PBS and protein binding

LogD_pH7.4_, stability and protein binding followed a previously described procedure ^70^. For stability and protein binding studies, samples were diluted to a concentration of 1.1 µM and 12 µM in the case of Gallium-68 and Zirconium-89 labelled probes.

### Tumour cell lines and cell culture

H2009 human lung adenocarcinoma cells were cultivated with DMEM:F12 medium (FG 4815, Biochrom, Berlin, Germany) supplemented with 5% (v/v) fetal bovine serum, FBS (10270, Invitrogen, Thermo Fisher Scientific, Waltham, Massachussetts, US), 1% (v/v) Penicillin Streptomicin Glutamin, PSG (10378, Gibco, Thermo Fisher Scientific, Waltham, Massachussetts, US), Insulin-Transferrin-Sodium Selenite (ITS) supplement (11074547001, Roche, Basel, Switzerland) 10 nM Hydrocortisone (H6909, Sigma-Aldrich, St. Louis, Missouri, US), 4.5 mM L-Glutamine (G7513, Sigma-Aldrich, St. Louis, Missouri, US), and 10 nM β-Estradiol (E2758, Sigma-Aldrich, St. Louis, Missouri, US). Cells were subcultured after trypsination in a ratio of 1:2–1:5, two to three times weekly at 70-80% confluency.

MDA-MB-231 breast cancer cells were cultivated with EMEM (41965039, Gibco, Thermo Fisher Scientific, Waltham, Massachussetts, US) supplemented with 10% (v/v) FBS, 1% (v/v) PSG and 1% (v/v) non-essential aminoacid (11140050, Gibco, Thermo Fisher Scientific, Waltham, Massachussetts, US).

Both cell lines were grown in a monolayer culture at 37°C in a 5% CO_2_ humidified atmosphere. Cell identity was authenticated, and cells were regularly tested for mycoplasma contamination.

### Cellular uptake assay

1.0 x 10^6^ cells per well were seeded in 6-well plates and grown for 2 days.

On the day of the experiment, cells were washed and then incubated in culturing medium with 1 nM/well of radioactive compound for 1h at 37°C. For blocking, the RGD alkyne peptide was added prior to the radiocompound to a final concentration of 1 µM/well. At the end of the incubation, the medium was removed and the cells rinsed with 2 x 1 mL of PBS/0.5% (w/v) Bovine Serum Albumin (BSA). Thereafter, they were washed twice with 1 mL of 50 mM glycine buffer (pH 2.8) with 0.1 M NaCl to remove the membrane-bound radiocompound. Finally, the cells were lysed with 2 x 1 mL of 1 M NaOH to determine the internalized radioligand. All fractions were measured in the γ-counter and the percentage of internalized and membrane bound radiocompound in relation to the total radioactivity added to the cells was reported.

### Integrin affinity evaluation

The integrin affinities of FSC(PEG4-αvβ6)_3_ and of [^nat^Ga]Ga-FSC(PEG4-αvβ6)_3_ were determined using an established ELISA protocol and expressed as 50% inhibitory concentrations (IC50)^67^. For ^nat^Ga-labelling, 10 µl (1 mM) of FSC(PEG4-αvβ6)_3_ were mixed with 65 µl of HEPES buffer (1.0 M) and 25 µl Ga(NO_3_)_3_ (2 mM) and heated to 50°C for 10 minutes. The solution was used without further purification.

### Animal experiments

All animal experiments were performed in accordance with the ethical standards of the institution in accordance with general animal welfare regulations in Austria and Germany and approved by the Austrian Ministry of Science or by the Regierung von Oberbayern. To assess biodistribution of ^68^Ga-FSC(PEG4-αvβ6)_3_ in healthy mice, we used 6-to 7-week-old female BALB/c mice (Charles River Laboratories, Wilmington, Massachusetts, US). For *in vivo* experiments in tumor-bearing animals, female CB17 severe combined immunodeficiency (SCID) mice were obtained from Charles River (Sulzfeld, Germany). At 6-10 weeks of age, mice were xenografted with 5 × 10^6^ H2009 cells in a 1:1 mixture of Medium:Matrigel® (Geltrex™ LDEV-Free Reduced Growth Factor Basement Membrane Matrix, A1413202, Life Technologies, Thermo Fisher Scientific). PET imaging and biodistribution studies were initiated when tumors had reached a diameter of approximately 7–10 mm (5–6 weeks after inoculation).

### Metabolic stability *in vivo*

One mouse was injected with 2.8 nmol of [^68^Ga]Ga-FSC(PEG4-αvβ6)_3_ (12.7 MBq) and sacrificed 15 min p.i. Urine and blood samples were collected at the time of sacrifice. The blood sample was centrifuged for 2 min at 18400 rcf. 100 µL of the supernatant was diluted and mixed 1:1 with ACN and centrifuged again to separate the protein pellet. An aliquot of the supernatant was diluted 1:1 with water and analysed via radio-RP-HPLC. Before the analysis the urine sample was solely diluted 1:100 with water.

### *Ex vivo* biodistribution in healthy mice

To evaluate *ex vivo* biodistribution, 3 healthy mice were injected with 0.15 nmol of [^68^Ga]Ga-FSC(PEG4-αvβ6)_3_ (0.5 MBq) and sacrificed after 90 min. The organs of interest were extracted, weighed and measured in the γ-counter. Results were expressed as percentage of injected dose per gram tissue (% ID/g).

### PET Imaging and biodistribution of tumor-bearing mice

We conducted PET imaging and biodistribution analysis in H2009 xenograft bearing mice to characterize the *in vivo* pharmacokinetic properties and tumor uptake of [^68^Ga]Ga-FSC(PEG4-αvβ6)_3_ and [^89^Zr]Zr-FSC(PEG4-αvβ6)_3_. To allow for head-to-head comparison, animals (n=3) were first imaged with [^68^Ga]Ga-Trivehexin (2-5 MBq, 53-68 pmol), followed 3 days later by [^68^Ga]Ga-FSC(PEG4-αvβ6)_3_ (4-7 MBq, 120-139 pmol). Imaging was carried out on a preclinical PET scanner (Nanoscan, Mediso), 75-90 min post i.v. tracer injection. Uptake specificity was assessed in a blocking group by pre-injection of cold labelled Trivehexin (50 nmol, 10 min prior to radiotracer) in n=2 animals. A separate cohort (n=3) also first received a benchmarking PET scan of [^68^Ga]Ga-Trivehexin (4-5 MBq, 50-80 pmol; 75-90 min p.i.), followed the day after by PET imaging after i.v. injection of [^89^Zr]Zr-FSC(PEG4-αvβ6)_3_ (0.4-0.6 MBq, 200-304 pmol; at 75 min, 24 h, 72 h, and 6 days p.i.;15 min per scan), followed by sacrifice and biodistribution. Additional animals (n=3/group) were injected for biodistribution analysis at 90 min, 90 min with blocking, and 24 h p.i.. PET data reconstruction, image analysis and quantification were performed using Nucline and Interview fusion software (both Mediso) without scatter and attenuation correction. PET reconstruction parameters: TT3D, It:4, Ss:6, 400–600 keV, 1:3, R:0.0005, M:24.

## Supporting information

Supporting Information FSC-integrin

## SUPPORTING INFORMATION

Supplementary material is available at… and includes detailed description of the instrumentation, of the synthesis of the labelling precursor and of additional experimental results.

## AUTHOR INFORMATION

## Author Contributions

G.G. synthesized the peptide conjugate, contributed to *in vitro* and *in vivo* characterization, performed data analysis and visualization, wrote the original draft, and edited the final manuscript. F.A.P. contributed to the *in vitro* characterization. M.A.Z. synthesized the peptide derivative. S.St. performed the ELISA-based affinity assays. N.H. and T.R. conducted the radiolabeling and quality controls for the *in vivo* experiments. S.St. conducted the xenografted animal experiments and S.K. analyzed the *in vivo* data. S.K. conceptualized the project and supervised the xenograft experiments. C.D. conceptualized and supervised the project and acquired funding. All authors reviewed and approved the final version of the manuscript.

## ACKNOWLEDGMENT

The authors gratefully acknowledge Christine Rangger for assistance with animal experiments at the Medical University Innsbruck, as well as Markus Mittelhäuser, Natalie Röder, and Sybille Reder for assistance with animal experiments at the Preclinical Imaging Core Facility (PICTUM) at TranslaTUM and TRIMT GmbH for providing Trivehexin.

