## Supporting Information FSC-integrin for "Synthesis and Preclinical Development of a Novel ^68^Ga/^89^Zr-Labelled ανβ6-Integrin Targeting Trimer"

### INSTRUMENTATION

#### Analytical RP-HPLC

RP-HPLC analysis was performed on a UltiMate 3000 system equipped with pump, autosampler, column compartment and diode array detector (Thermo Fisher Scientific, Vienna, Austria).

Method A(A): A Dr. Maisch ReproSil Pur C18 AQ, 120 Å, 5 µm, 150 x 4.6 mm (Dr. Maisch GmbH Ammerbuch-Entringen, Germany) column with a flow rate of 1 mL/min and UV detection at 220 nm was used. Acetonitrile (ACN)/$H_2O$ + 0.1% trifluoroacetic acid (TFA) was used as mobile phase with the following multistep gradient: 0.0-1.0 min 25% ACN, 1.0-16.0 min 25-40% ACN, 16.0-17.0 min 40-95% ACN, 17.0-21.0 min 95% ACN, 21.0-21.5 min 95-25% ACN, 21.5-24.0 min 25% ACN.

Method A(B): A Jupiter 4 µm Proteo 90 Å 250 x 4.6 mm (Phenomenex Ltd. Aschaffenburg, Germany) column with flow rate of 1 mL/min and UV detection at 220 nm was used. Acetonitrile (ACN)/$H_2O$ + 0.1% trifluoroacetic acid (TFA) was used as mobile phase with the following multistep gradient: 0.0-3.0 min 10% ACN, 3.0-16.0 min 10-60% ACN, 16.0-18.0 min 60% ACN, 18.0-18.1 min 60-10% ACN, 18.1-22.0 min 10% ACN.

#### Preparative RP-HPLC

RP-HPLC purification was performed on a UltiMate 3000 pump with UltiMate 3000 UV/Vis detector (Thermo Fisher Scientific, Vienna, Austria).

Method P(A): A Nucleodur 7 µm C18 HTec 250 x 21 mm (Macherey-Nagel, Düren, Germany) column with flow rate of 15 mL/min and UV detection at 440 nm was used. Acetonitrile

(ACN)/$H_2O$ + 0.1% trifluoroacetic acid (TFA) was used as mobile phase with the following multistep gradient: 0.0-5.0 min 3% ACN, 5.0-7.0 min 3-12% ACN, 7.0-17.0 min 12-22% ACN, 17.0-17.5 min 22-3% ACN, 17.5-27.0 min 3% ACN.

Method P(B): A Nucleodur 5 µm C18 HTec 250 x 16 mm (Macherey-Nagel, Düren, Germany) column with flow rate of 8 mL/min and UV detection at 220 nm was used. Acetonitrile (ACN)/$H_2O$ + 0.1% trifluoroacetic acid (TFA) was used as mobile phase with the following multistep gradient: 0.0-5.0 min 10% ACN, 5.0-25.0 min 10-65% ACN, 25.0-25.1 min 65-10% ACN, 25.1-34.0 min 10% ACN.

Method P(C): A Nucleodur 5 µm C18 HTec 250 x 16 mm (Macherey-Nagel, Düren, Germany) column with flow rate of 8 mL/min and UV detection at 220 nm was used. Acetonitrile (ACN)/$H_2O$ + 0.1% trifluoroacetic acid (TFA) was used as mobile phase with the following multistep gradient: 0.0-5.0 min 4% ACN, 5.0-37.0 min 4-55% ACN, 37.0-37.1 min 55-4% ACN, 37.1-46.0 min 4% ACN.

**ESI-MS**

All ESI-MS experiments were carried out with LCMS-2050 Nexera (Shimadzu, Kyoto, Japan) using the following conditions: scan range (m/z 200.0 – 2000; sampling 500 msec /2Hz; mobile phase 30/70 H2O/ACN +0.1 % Formic Acid; flow rate of 0.5 mL/min). Data were acquired and evaluated with LabSolutions software (Shimadzu, Kyoto, Japan).

**MATERIALS**

All commercially available chemicals, reagents and solvents were of analytical grade and were used without further purification. Only high-purity water (18 mΩ) was employed. Glycine-2-

Chlorotrityl resin (H-Gly-2-ClTrt) was purchased from Novabiochem (Merck KGaA, Darmstadt, Germany). Coupling reagent 1-hydroxy-7-azabenzotriazole (HOAt) 1 M solution in DMA was obtained from Activate Scientific GmbH (Prien, Germany). Azido-PEG4-acid was purchased from BroadPharm (San Diego, CA, USA). N-Methyl-2-pyrrolidon (NMP) was obtained from Carl Roth (Karlsruhe, Germany). All other reagents were purchased from Sigma-Aldrich (Merck KGaA, Darmstadt, Germany) or Merck (Darmstadt, Germany).

### METHODS

#### Synthesis of the RGD alkyne derivative

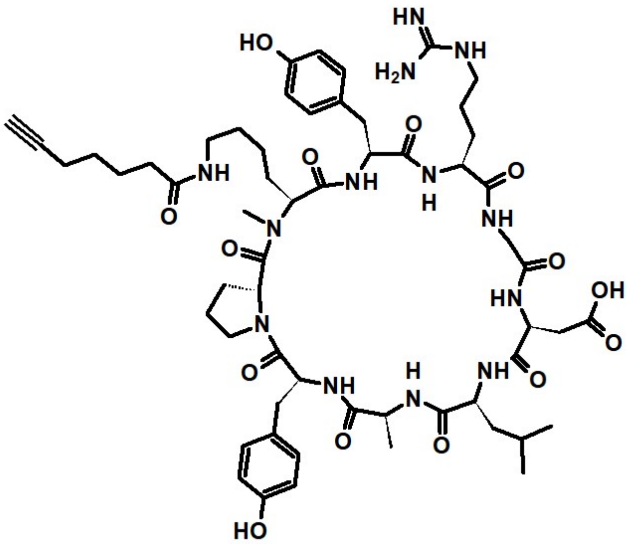

**Figure S1:** Chemical structure of the RGD alkyne derivative: cyclo[YRGDLAYp(*N*Me)K(6-heptynoic amide)].

General procedures (GP) for solid-phase peptide chemistry

*On-resin peptide bond formation for single amino acids (GP1)*

In a 5 mL glass vial HOAt (2.0 eq), HATU (2.0 eq), and the respective amino acid (2.0 eq) were dissolved in 4 mL DMF with stirring. DIPEA (6.0 eq) was added and the solution was stirred for another 5 min at room temperature. This mixture was added to the H-Gly-2-ClTrt resin (1.0 eq) and the reaction was allowed to proceed for 3 hours. Afterward, the resin was washed with DMF (3 × 6 mL/g resin).

*Fmoc-removal (GP2)*

The resin was treated with a mixture of 20 % Piperidine/DMF (vol/vol) (1 × 5 min, 1 × 15 min), followed by a washing step with DMF. (8 × 6 mL/g resin).

*On-resin Dde-deprotection* (*GP3*)

For Dde-deprotection in the presence of Fmoc-groups, the resin was treated with a solution of Imidazole (0.92 g/g resin) and Hydroxylamine Hydrochloride (1.26 g/g resin) in 5 mL of NMP and 1 mL of DMF. After 3 hours of incubation, the resin was washed with DMF (3 × 6 mL/g resin).

*N-methylation (GP4)*

Prior to proceeding with the *N*-methylation step, the resin was thoroughly washed with DCM (3 × 6 mL/g of resin) to ensure complete removal of DMF and to avoid otherwise the formation of a reddish, gel-like precipitate in the subsequent reaction step.

First, an adjuvant *N*-protecting group was introduced. Therefore, 1-(chlorosulfonyl)-2-nitrobenzene (4.0 eq) was dissolved in 2,4,6-collidine (10.0 eq) and DCM (6 mL/g resin). This solution was added to the resin for 15 min, followed by a washing step with DCM (3 × 6 mL/g resin) and THF (3 × 6 mL/g resin). For the *Mitsunobu* reaction, Triphenylphosphine (5.0 eq) was dissolved in 2 mL of dry THF and 10.0 eq Methanol was added. The resin was immersed in this solution for 2 min. To this, a second solution of Diisopropylazidodicarboxylate (5.0 eq) in 3 mL dry THF was slowly added with caution. After 10 min of incubation, the resin was washed with THF (3 × 6 mL/g resin). Removal of 1-(chlorosulfonyl)-2-nitrobenzene occurred within 5 min by treating the resin with a mixture of mercaptoethanol (10.0 eq) and 1,8-Diazabicyclo[5.4.0]undec-7-ene (5.0 eq). The resin was washed with DMF (8 × 6 mL/g resin) afterwards.

*Capping of unreacted amines (GP5)*

The resin was immersed in 5 mL of a freshly prepared Acetic Anhydride/Pyridine (3:2) solution for 30 min. Afterwards, it was washed three times with DMF.

*Cleavage of protected peptides from the resin (GP6)*

The resin was washed thoroughly with DCM (3 × 6 mL) before 5 mL of a cleavage cocktail containing 1,1,1,3,3,3-hexafluoroisopropanol/DCM (80/20) (vol/vol) was added. After 45 min of incubation, the crude peptide was transferred into a 500 mL round bottom flask and 5 mL of fresh cleavage cocktail was added to the syringe. This step was repeated three times in total. The resin was washed with DCM and the wash fractions were also added to the round bottom flask. All volatiles were reduced *in vacuo*.

*Cyclization (GP7)*

The peptide was dissolved in 250 mL DMF with stirring and 3.0 eq of Diphenylphosphorylazide as well as 5.0 eq of solid $NaHCO_3$. After 24 hours, the reaction mixture was concentrated *in vacuo* and filtered, followed by a complete removal of all volatiles *in vacuo.*

Synthesis of cyclo[YRGDLAYp(*N*Me)K(6-heptynoic amide)] peptide

For the synthesis of the cyclic peptide 500 mg of a H-Gly-2-ClTrt resin (δ = 1.1 mmol/g) was weighed into a 20 mL plastic syringe, equipped with a pp-frit inlet, and the resin was swollen for 1 hour in DMF. Coupling of Fmoc-L-Arg(Pbf)-OH, Fmoc-L-Tyr(tBu)-OH and Fmoc-L-

Lys(Dde)-OH followed *GP1* & *GP2*. On-resin Dde-removal was performed according to *GP3*. Attachment of the alkyne tag to the lysine sidechain required only 1.2 eq HOAt, 1.2 eq HATU, 1.2 eq 6-heptynoic acid and 3.0 eq DIPEA. After the Fmoc-deprotection of Lysine (*GP2*), the *N*-methylation was carried out (*GP4*). The coupling of Fmoc-D-Pro-OH (*GP1* & *GP2*), was accompanied by a capping step *(GP5*) to avoid the formation of truncated peptide species. The assembly of the remaining amino acids (Fmoc-L-Tyr(tBu)-OH, Fmoc-L-Ala-OH, Fmoc-L-Leu-OH and Fmoc-L-Asp(tBu)-OH) followed *GP1* & *GP2* as usual. The functionalized nonapeptide was cleaved from the solid support according to *GP6* and directly cyclized (*GP7*). Purification *via* semipreparative HPLC (gradient A(A)) and subsequent lyophylization yielded 23.2 mg (19.6 µmol, 3.5 %) of a colorless solid with a purity > 98% confirmed by analytical RP-HPLC (gradient A(A); $t_R$ = 9.9 min). ESI/MS: *m/z* [M+H$^+$] = 1186.0 [$C_{58}H_{84}N_{13}O_{14}^+$], exact mass (monoisotopic): 1186.6 (calculated), *m/z* [M+2 H$^+$] = 593.9 [$C_{58}H_{85}N_{13}O_{14}^{2+}$], exact mass (monoisotopic): 593.8 (calculated).

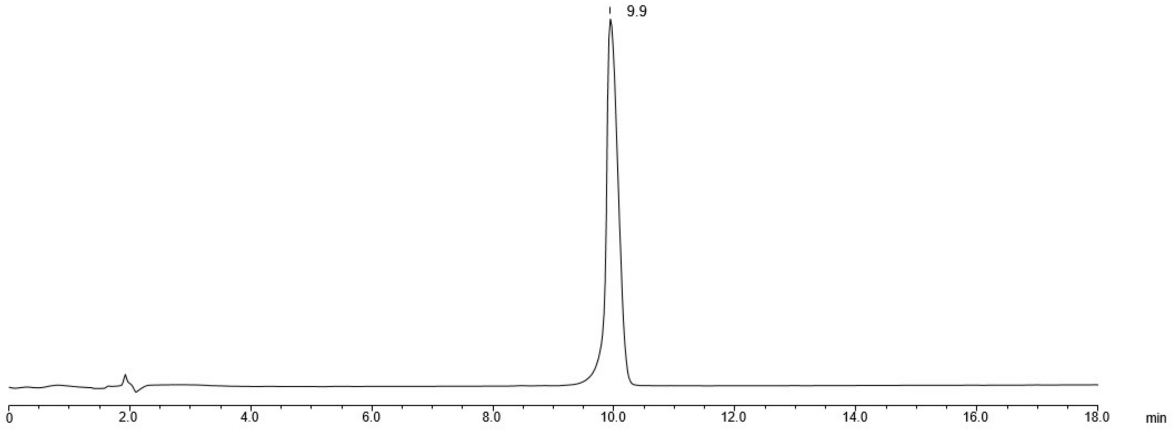

**Figure S2:** UV-HPLC chromatogram of the RGD alkyne derivative.

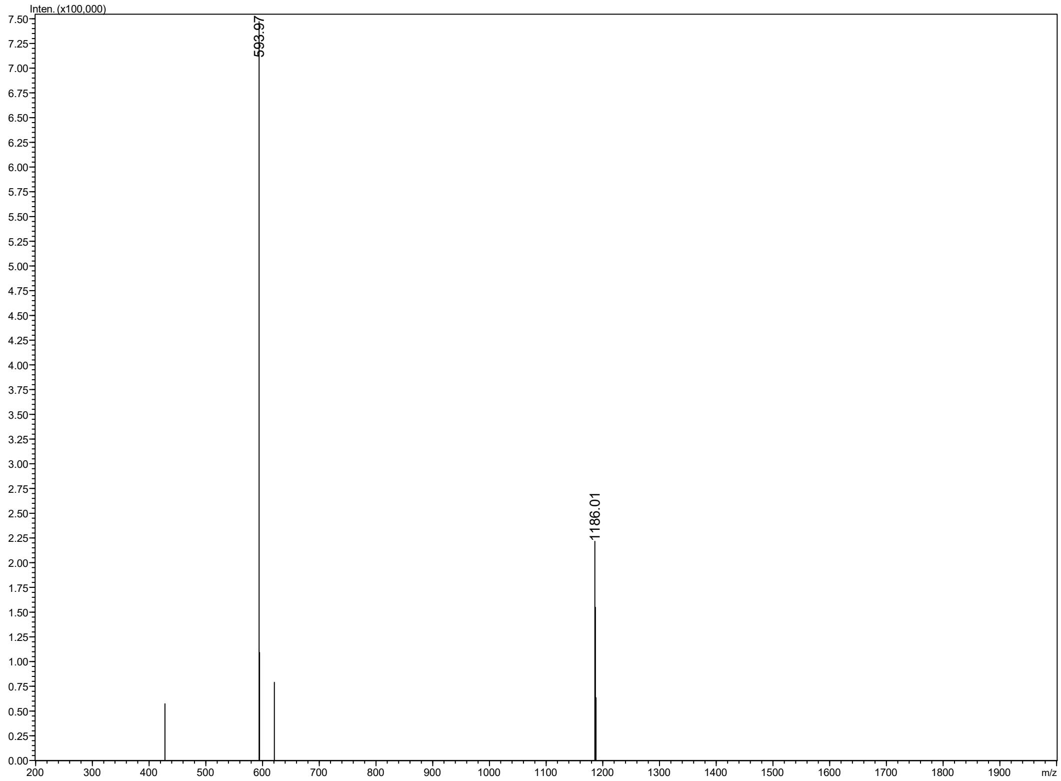

**Figure S3:** MS spectrum of the RGD alkyne derivative.

### Synthesis of the FSC(PEG4-αvβ6)$_3$

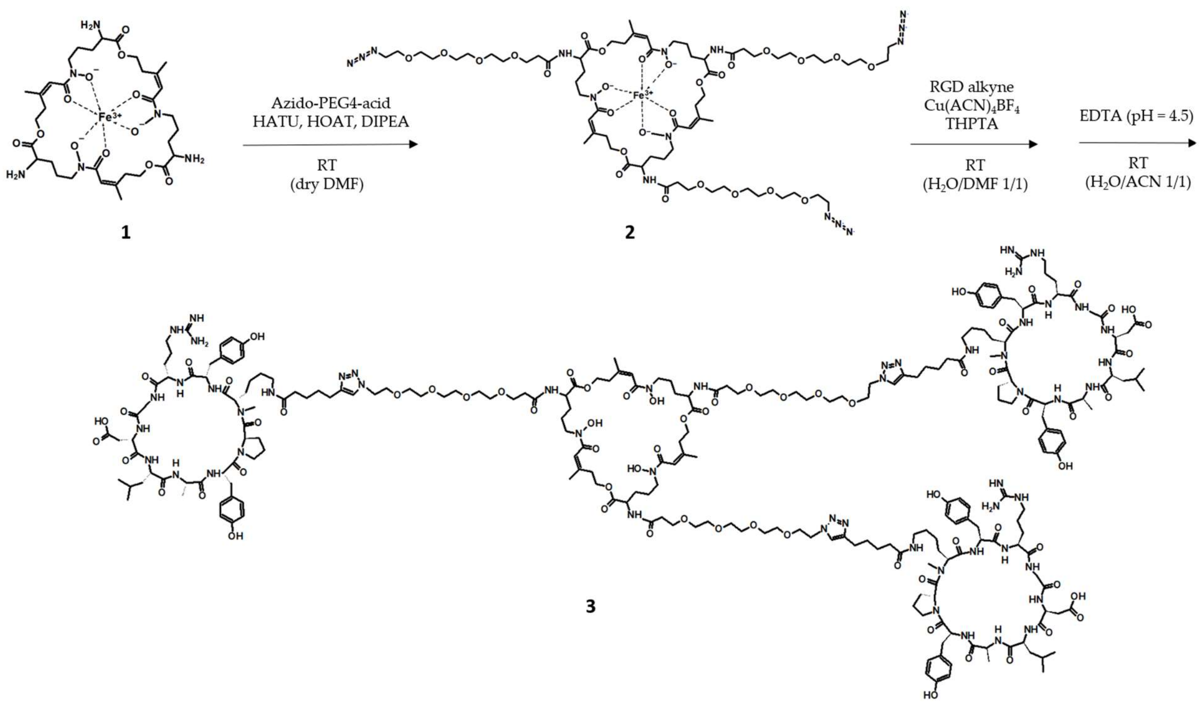

**Figure S4:** Synthetic scheme of the FSC(PEG4-αvβ6)$_3$ ligand.

<u>Extraction of Fusarinine C (FSC) (compound 1 in Fig. S4)</u>

The siderophore-based natural product fusarinine C (FSC) (**1**) was isolated from 5 liters of Aspergillus fumigatus ΔsidG culture, grown under iron-deficient conditions as described by Schrettl and co-workers[1]. Following filtration of the culture supernatant, an excess of $FeSO_4$ or $FeCl_3$ was added to a final concentration of 10 mM, resulting in a red-coloured solution indicating the formation of the complex. The sediment present in the solution was separated by centrifugation (5 min at 3220 x g and 25°C).

After preparative RP-HPLC purification (gradient P(A); $t_R$ = 14.2 min) and freeze-drying, 735 mg of a red-brown powder was obtained with a purity > 94 % confirmed by analytical RP-HPLC (gradient A(B); $t_R$ = 9.3 min). ESI/MS: *m/z* [M+H$^+$] = 780.2 [$C_{33}H_{52}FeN_6O_{12}^+$], exact mass (monoisotopic): 780.3 (calculated), *m/z* [M+2 $H^+$] = 390.8 [$C_{33}H_{53}FeN_6O_{12}^{2+}$], exact mass (monoisotopic): 390.6 (calculated).

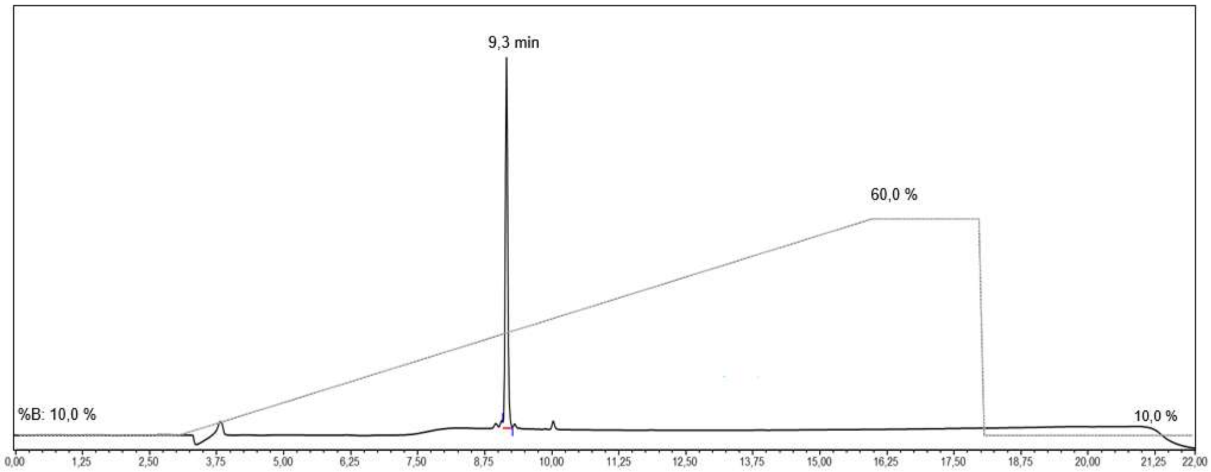

**Figure S5:** UV-HPLC chromatogram of [Fe]FSC.

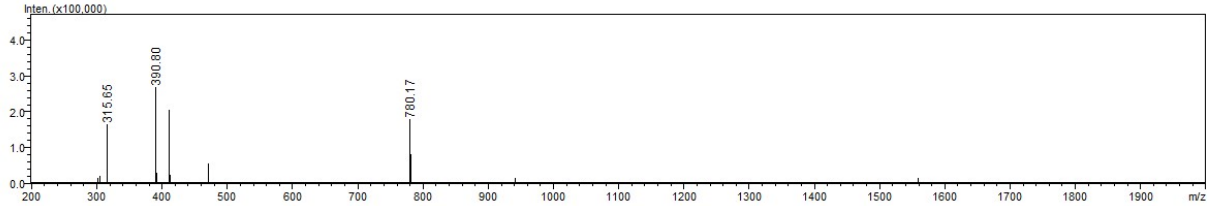

**Figure S6:** MS spectrum of [Fe]FSC.

[Fe]FSC derivatisation_[Fe]FSC(PEG4-Azido)$_3$ (compound 2 in Fig. S4)

4 mg of the siderophore [Fe]FSC (**1**) (5.1 µmol, 1.0 eq.) were dissolved in 125 µL of dry DMF and the pH adjusted to 9 with DIPEA.

29.3 mg of HATU (76.9 µmol), 10.5 mg of HOAt (76.9 µmol) were dissolved with 814 µL of dry DMF and then mixed with 7.5 mg of Azido-PEG4-Acid (25.7 µmol, 5.0 eq.). The pH was corrected to 9 with 23 µL of DIPEA and the resulting solution was allowed to rest for approximately 5 min. Thereafter, it was added dropwise to the solution of the siderophore and the final pH checked and adjusted to 9 if necessary. After 30 min the reaction was stopped by diluting the solution 1:1 with water.

Preparative RP-HPLC purification (gradient P(B); $t_R$ = 26.1 min) yelded 4.7 mg (2.9 µmol, 56.9 %) of [Fe]FSC(PEG4-Azido)$_3$ with a purity > 99 % confirmed by analytical RP-HPLC ($t_R$ = 17.4 min). ESI/MS: *m/z* [M+H$^+$] = 1599.5 [$C_{66}H_{109}FeN_{15}O_{27}^{+}$], exact mass (monoisotopic): 1599.7 (calculated), *m/z* [M+2 H$^+$] = 800.5 [$C_{66}H_{110}FeN_{15}O_{27}^{2+}$], exact mass (monoisotopic): 800.4 (calculated).

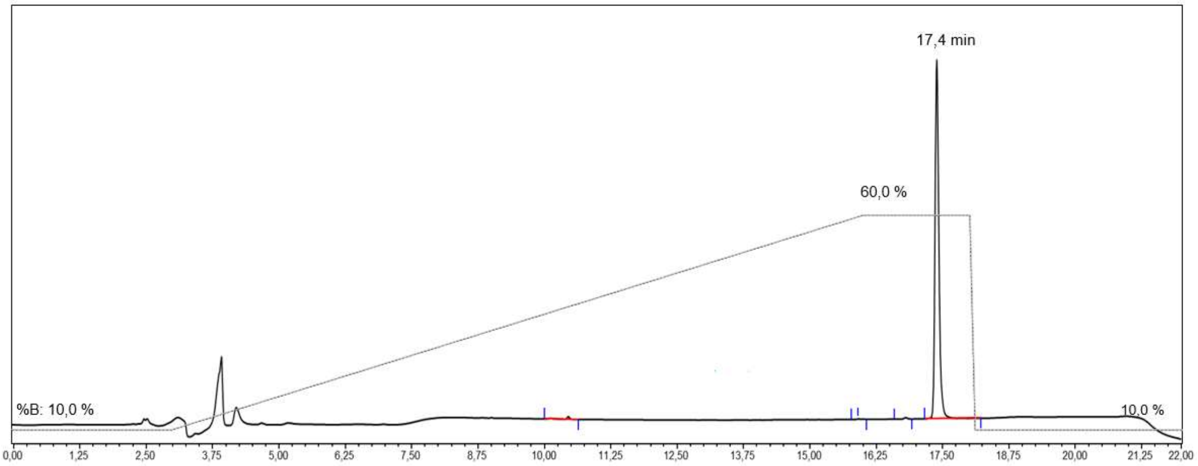

**Figure S7:** UV-HPLC chromatogram of [Fe]FSC(PEG4-Azido)$_3$.

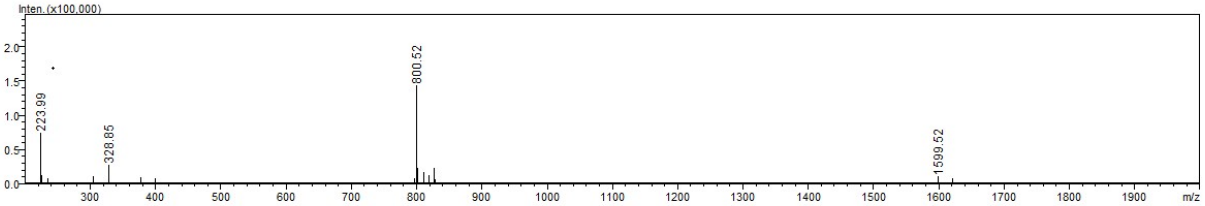

**Figure S8:** MS spectrum of [Fe]FSC(PEG4-Azido)$_3$.

<u>FSC(PEG4-αvβ6)$_3$ synthesis (compound 3 in Fig. S4)</u>

4.7 mg of [Fe]FSC(PEG4-Azido)$_3$ (**2**) (2.9 µmol, 1.0 eq.) and 13.9 mg of RGD alkyne (11,8 µmol, 4.0 eq.) was dissolved with 550 µL of $H_2O$/DMF 1/1 and degassed with Ar(g) flow for 10 min. 12.8 mg of THPTA (29.4 µmol, 10.0 eq.) and 9.2 mg of $Cu(ACN)_4BF_4$ catalyst (29.4 µmol, 10.0 eq.) were dissolved separately in dry DMF and degassed with Ar(g) flow for 15 min, then mixed together to give a clear green solution and degassed for further 5 min. Eventually the catalyst solution was added to the solution of azide and alkyne and let react under Ar(g) atmosphere for 1 h.

For demetallation, the organic solvent was evaporated. The resulting conjugate was dissolved in 1 mL of $H_2O$/ACN 1/1 solvent and an aqueous solution of EDTA (400 mM) was added to provide an excess of approximately 50 eq. of EDTA over the conjugate. The pH was adjusted to 4.5 and the solution was stirred for 4 hours.

The crude mixture was dried by rotovap, redissolved in 1 mL of $H_2O$ + 30% ACN and then purified by preparative RP-HPLC (gradient P(C); tR = 30,1 min), yelding 2.0 mg (0.39 µmol, 13.4 %) of FSC(PEG4-αvβ6)$_3$ with a purity of > 97% confirmed by analytical RP-HPLC ( $t_R$ = 13.7 min). ESI/MS: *m/z* [M+4 $H^+$] = 1276.9 [$C_{240}H_{364}N_{54}O_{69}{}^{4+}$], exact mass (monoisotopic): 1276.7 (calculated), *m/z* [M+5 $H^+$] = 1021.5 [$C_{240}H_{365}N_{54}O_{69}{}^{5+}$], exact mass (monoisotopic): 1021.5 (calculated).

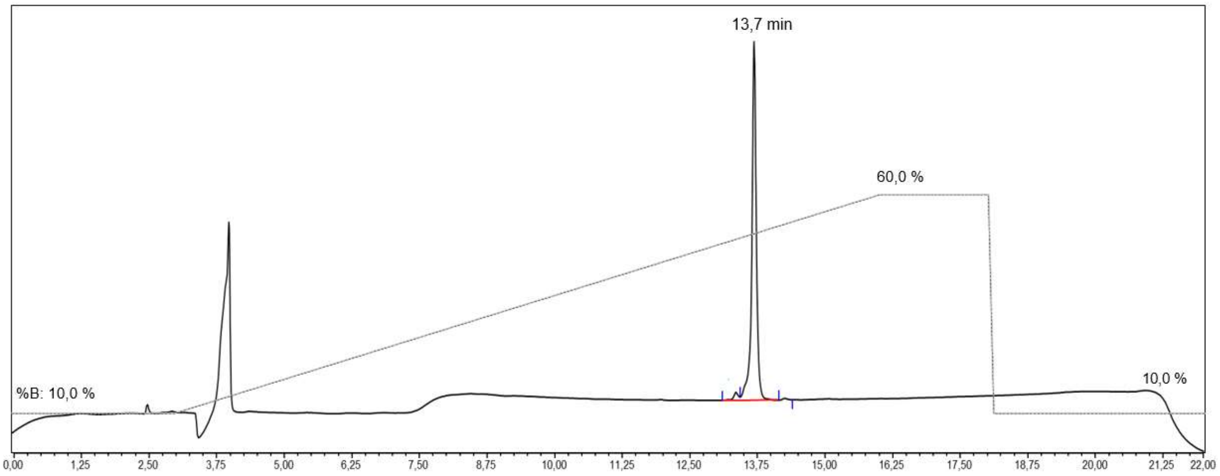

**Figure S9:** UV-HPLC chromatogram of FSC(PEG4-αvβ6)$_3$.

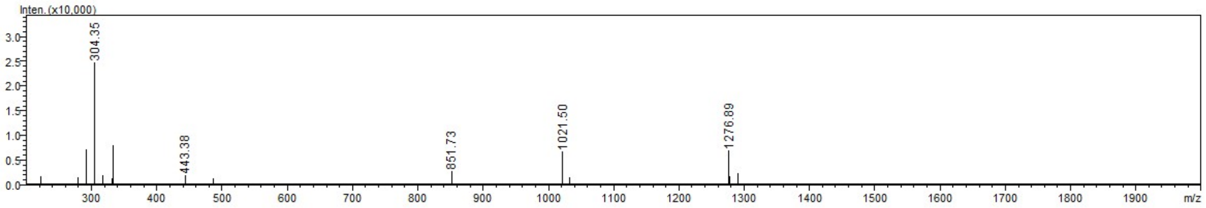

**Figure S10:** MS spectrum of FSC(PEG4-αvβ6)$_3$.

### ADDITIONAL RESULTS

#### Radiolabelling and quality control results by *radio*-HPLC and *radio*-iTLC

<u>Radiolabelling for [$^{68}$Ga]Ga/[$^{89}$Zr]Zr-FSC(PEG4-αvβ6)$_3$ and for [$^{68}$Ga]Ga-Trivehexin</u>

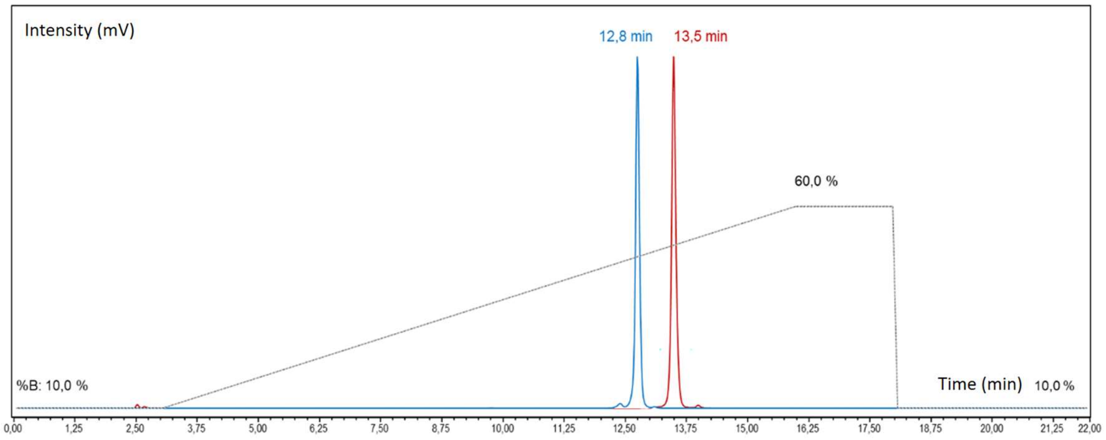

A

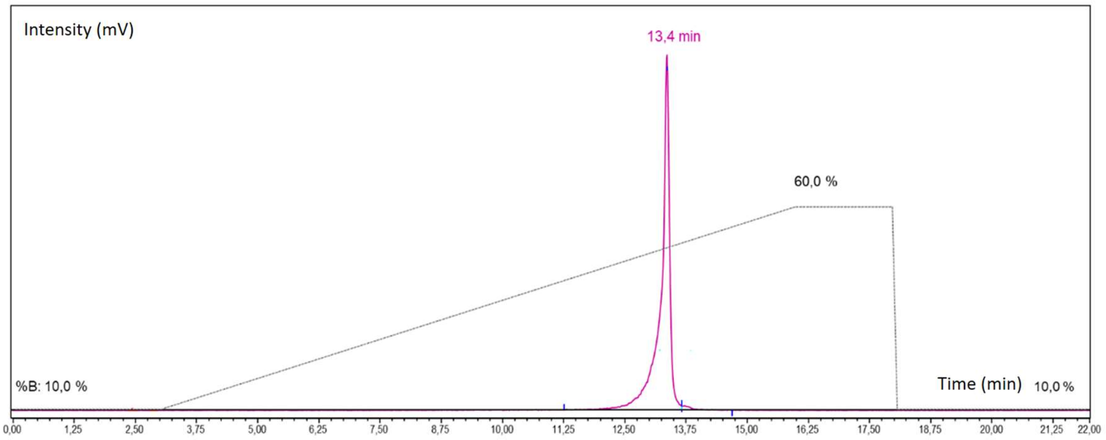

B

**Figure S11:** Radio RP-HPLC of [$^{68}$Ga]Ga-FSC(PEG4-αvβ6)$_3$ (red) and [$^{68}$Ga]Ga-Trivehexin (blue) (**A**) and of [$^{89}$Zr]Zr-FSC(PEG4-αvβ6)$_3$ (magenta) (**B**). Acetonitrile (ACN)/$H_2O$ + 0.1% trifluoroacetic acid (TFA) was used as mobile phase with the following multistep gradient: 0.0-3.0 min 10% ACN, 3.0-16.0 min 10-60% ACN, 16.0-18.0 min 60% ACN, 18.0-18.1 min 60-10% ACN, 18.1-22.0 min 10% ACN.

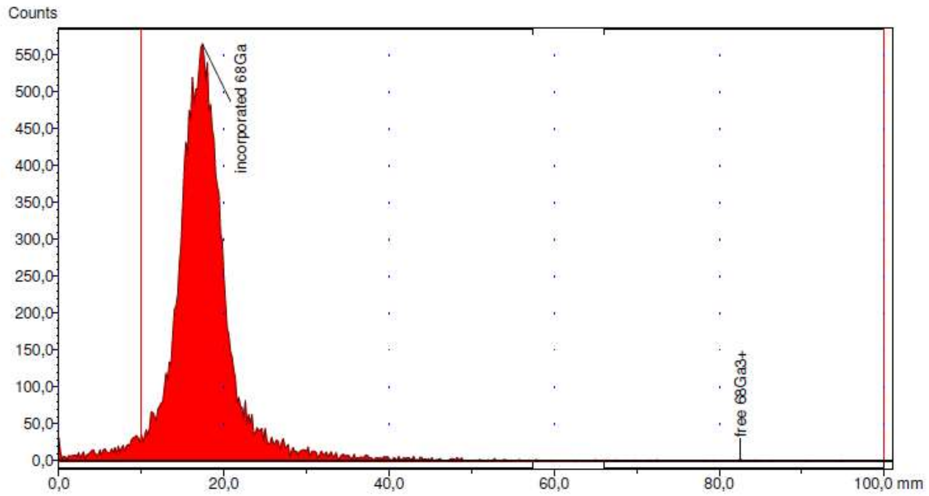

A

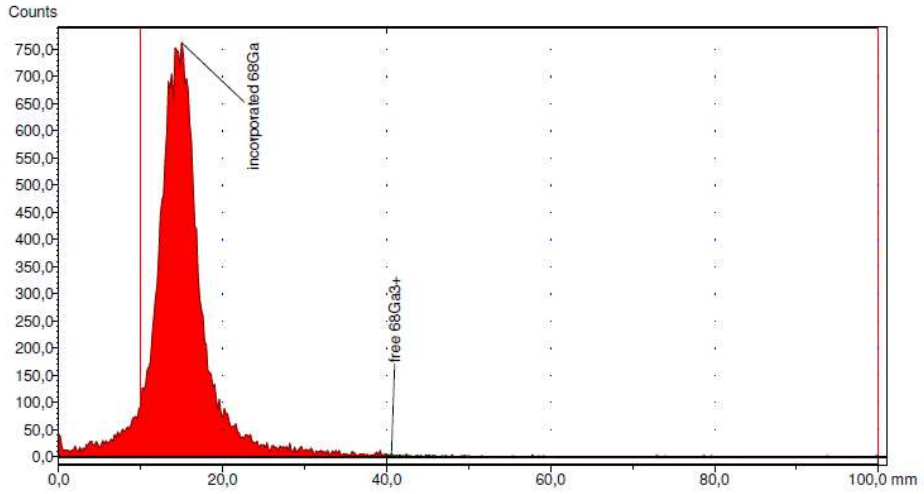

B

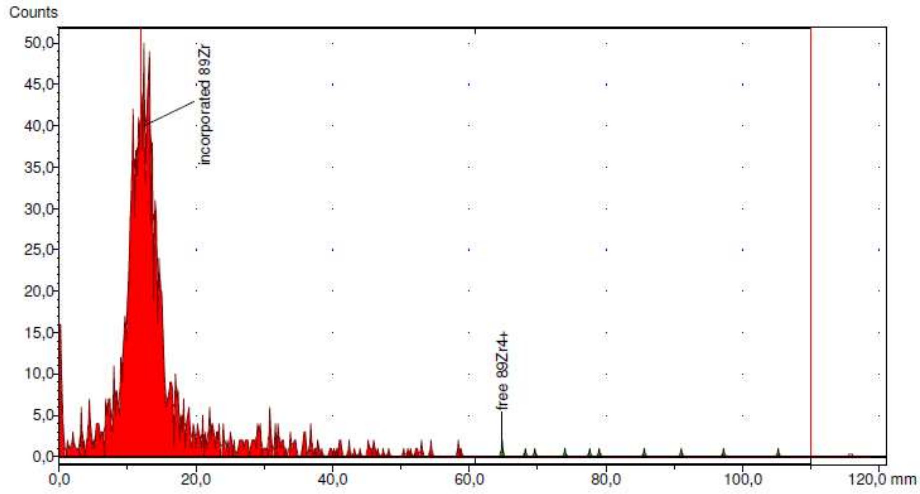

C

**Figure S12:** Representative *radio*-iTCL scans of [[68]Ga]Ga-FSC(PEG4-αvβ6)$_3$ (**A**), of [[68]Ga]Ga-Trivehexin (**B**) and of [[89]Zr]Zr-FSC(PEG4-αvβ6)$_3$ (**C**).

Zirconium-89 labelling of FSC(PEG4-αvβ6)$_3$ at varying molar activities

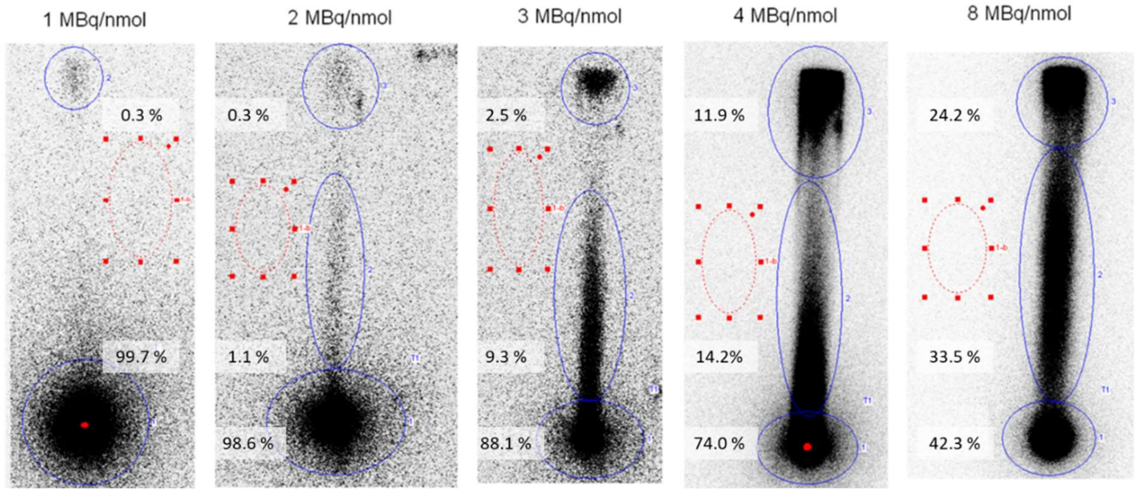

**Figure S13:** *Radio*-iTLC scans of the strips spotted with the labelling solution and eluted with 0.05 M EDTA solution (pH 7).Blue circles indicate the ROIs for the three quantified phases per run, red circles indicate the ROI used for background measurement.

**Results of the *in vitro* characterization**

Determination of the ligand affinity to the αvβ6 integrin

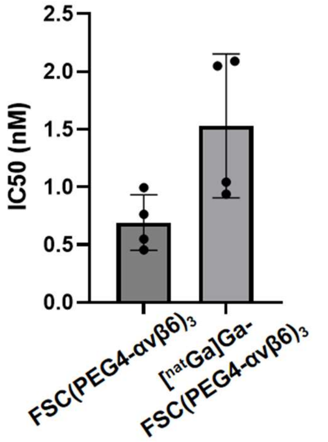

**Figure S14:** Affinity determination (IC50) of the unlabelled FSC(PEG4-αvβ6)$_3$ and of the [$^{nat}$Ga]Ga-FSC(PEG4-αvβ6)$_3$ obtained with four technical replicates.

Lipophilicity results for [$^{68}$Ga]Ga/[$^{89}$Zr]Zr-FSC(PEG4-αvβ6)$_3$

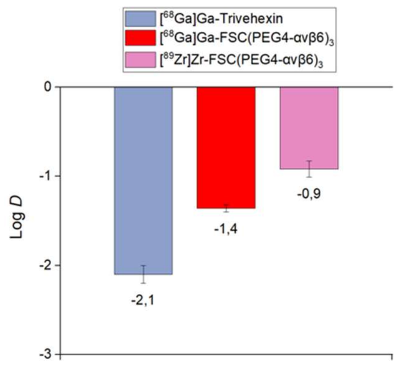

**Figure S15:** Distribution coefficients (LogD$_{pH7.4}$) of the Gallium-68 and Zirconium-89 labelled compounds obtained with six and five technical replicates, respectively. For comparison, the literature-reported LogD$_{pH7.4}$ value of [$^{68}$Ga]Ga-Trivehexin was also plotted[2].

Protein binding results for [$^{68}$Ga]Ga/[$^{89}$Zr]Zr-FSC(PEG4-αvβ6)$_3$ and for [$^{68}$Ga]Ga-Trivehexin

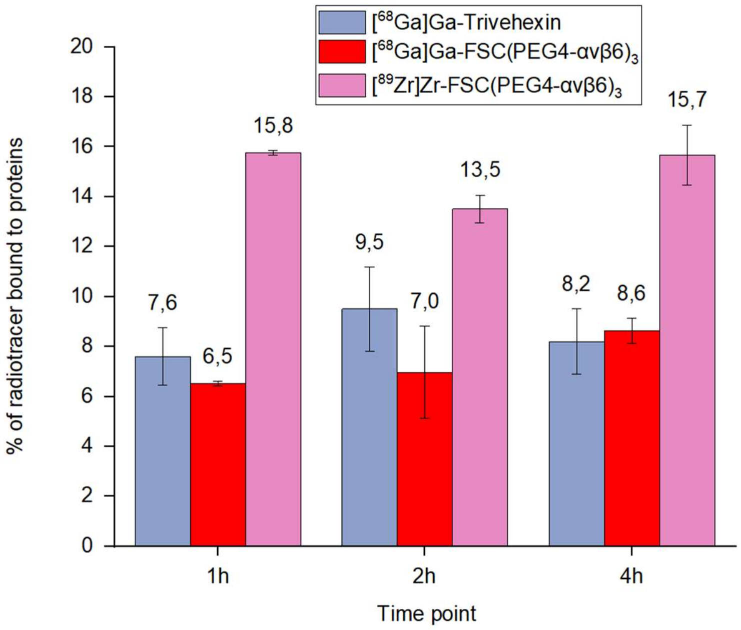

**Figure S16:** Binding to human serum proteins for Gallium-68 and the Zirconium-89 labelled FSC(PEG4-αvβ6)$_3$ and for [$^{68}$Ga]Ga-Trivehexin. Two independent experiments were conducted, each including two technical replicates.

Results of the *in vitro* stability investigation for [$^{68}$Ga]Ga/[$^{89}$Zr]Zr-FSC(PEG4-αvβ6)$_3$

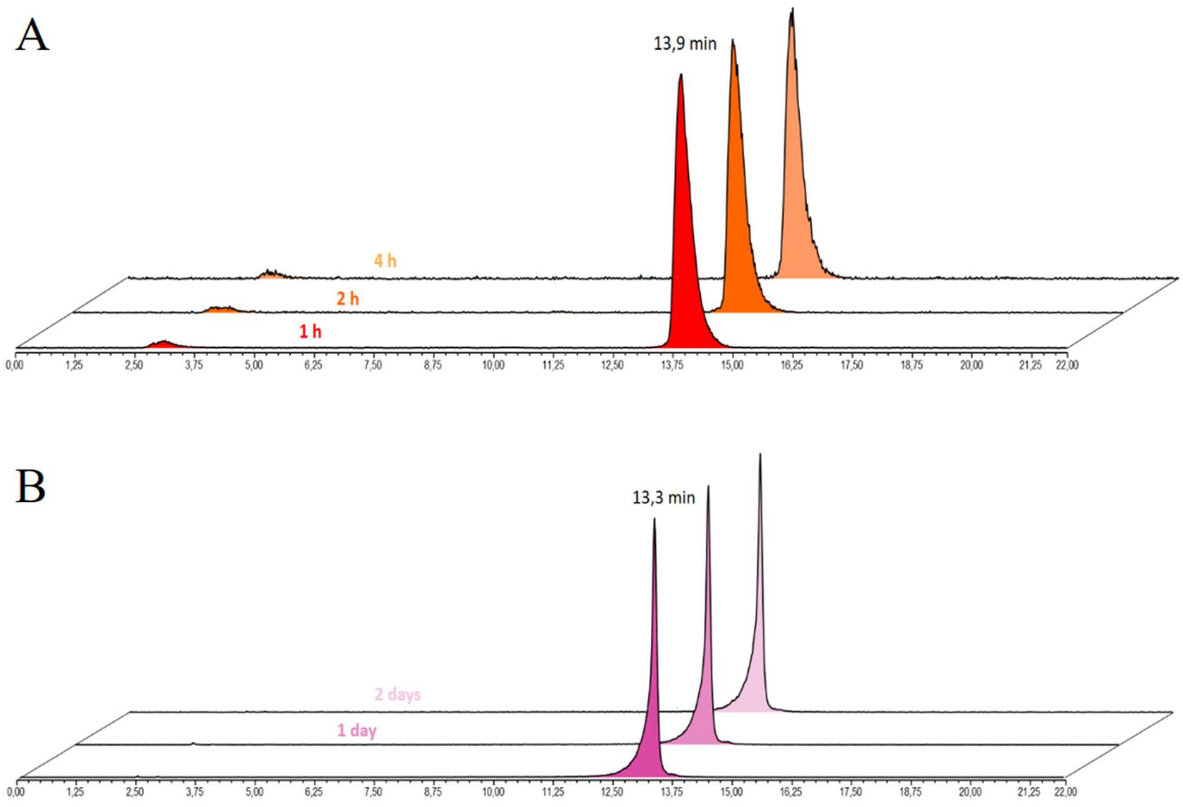

**Figure S17:** Radio-HPLC chromatograms for [$^{68}$Ga]Ga-FSC(PEG4-αvβ6)$_3$ in PBS (**A**) and for [$^{89}$Zr]Zr-FSC(PEG4-αvβ6)$_3$ labelling solution (**B**) at different timepoints.

**Ex vivo biodistribution results for [$^{68}$Ga]Ga-FSC(PEG4-αvβ6)$_3$ in healthy animals**

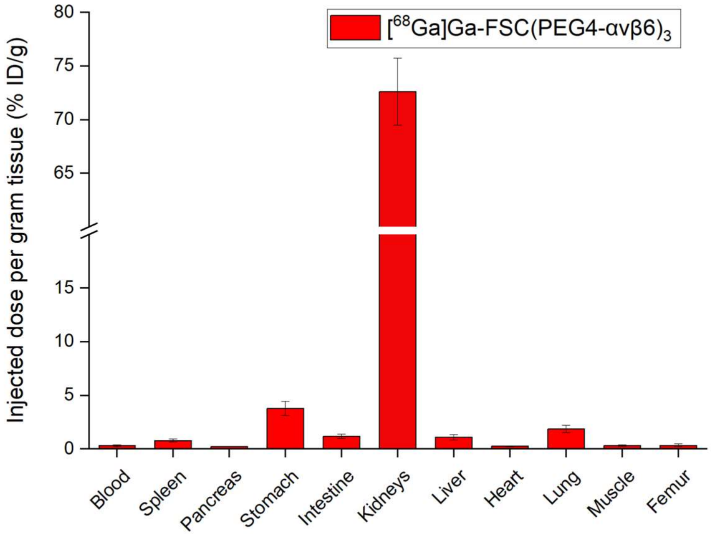

**Figure S18:** Ex vivo biodistribution study in healthy BALB/C mice (n = 3), performed 90 min p.i. for [$^{68}$Ga]Ga-FSC(PEG4-αvβ6)$_3$ (amount injected 0.15 nmol, 0.5 MBq).

**Metabolic stability for [$^{68}$Ga]Ga-FSC(PEG4-αvβ6)$_3$**

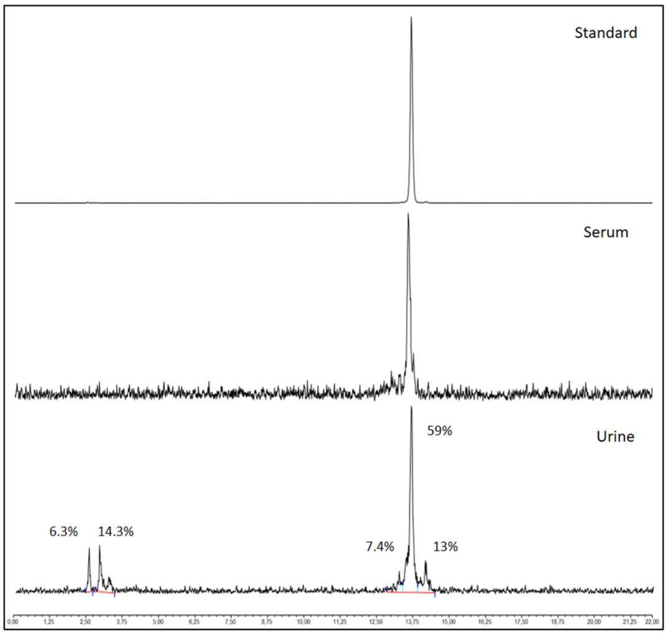

**Figure S19:** Radio-HPLC chromatograms for [$^{68}$Ga]Ga-FSC(PEG4-αvβ6)$_3$ injection solution (2.8 nmol,12.7 MBq) in mouse serum and urine 15 min p.i.

#### *Ex vivo* biodistribution comparison of [$^{68}$Ga]Ga-FSC(PEG4-αvβ6)$_3$ and [$^{68}$Ga]Ga-Trivehexin

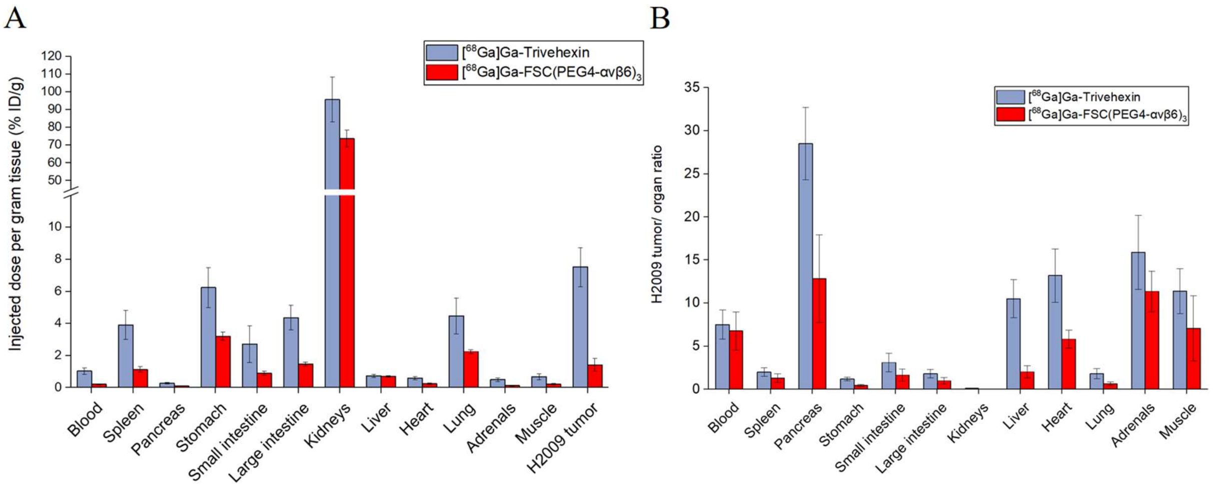

**Figure S20:** *Ex vivo* biodistribution study in H2009-bearing SCID mice (n = 3) performed 90 min p.i. for [$^{68}$Ga]Ga-FSC(PEG4-αvβ6)$_3$ (amount injected 120-140 pmol, 6.8-8.0 MBq). Literature values of [$^{68}$Ga]Ga-Trivehexin biodistribution at 90 min p.i. (n =5; amount injected 97 ± 13 pmol) obtained in the same model are plotted for comparison[2]. (**B**) Tumor-to-organ ratios derived from H2009 biodistribution data for [$^{68}$Ga]Ga-FSC(PEG4-αvβ6)$_3$ and [$^{68}$Ga]Ga-Trivehexin as reported in the literature[2].

(1) Schrettl, M.; Bignell, E.; Kragl, C.; Sabiha, Y.; Loss, O.; Eisendle, M.; Wallner, A.; Arst Jr, H. N.; Haynes, K.; Haas, H. Distinct roles for intra-and extracellular siderophores during Aspergillus fumigatus infection. *PLoS pathogens* **2007**, *3* (9), e128.
(2) Quigley, N. G.; Steiger, K.; Hoberück, S.; Czech, N.; Zierke, M. A.; Kossatz, S.; Pretze, M.; Richter, F.; Weichert, W.; Pox, C. PET/CT imaging of head-and-neck and pancreatic cancer in humans by targeting the "Cancer Integrin" αvβ6 with Ga-68-Trivehexin. *European Journal of Nuclear Medicine and Molecular Imaging* **2022**, 1-12.
